# Morphofunctional evaluation of the adrenal gland in rats submitted to nutritional restriction during pregnancy

**DOI:** 10.1101/2022.10.16.512413

**Authors:** Bruno dos Santos Telles, Hércules Jonas Rebelato, Marcelo Augusto Marretto Esquisatto, Rosana Catisti

**Affiliations:** Programa de Pós-graduação em Ciências Biomédicas, Centro Universitário da Fundação Hermínio Ometto – FHO, Araras, São Paulo, Brasil

**Keywords:** caloric restriction, protein restriction, pregnancy, adrenal, glucocorticoid/mineralocorticoid receptors.

## Abstract

Poor nutrition during pregnancy causes permanent metabolic and/or structural adaptation in offspring. The adrenal gland produces various steroid hormones during pregnancy. Thus, this study aimed to evaluate the influence of diet during pregnancy on the adrenal glands of Wistar rats. For this, 10-week-old pregnant Wistar rats (p, n=15) and non-pregnant rats (np, n=15) were divided into three groups and received a normoproteic control diet (C, 17% casein, n=5), isocaloric low-protein diet (PR, 6% casein, n=5), or 50% calorie restriction (CR, 50% of the diet consumed by group C), over a period of 21 days. On the 21st day of gestation (21dG, p groups) or on the 21st day of diet (np groups), after anesthetic deepening, the right adrenal gland was collected, weighed (total mass), and prepared for inclusion in Paraplast® for histomorphometric and immunohistochemical analysis (Ki-67, glucocorticoid receptors (GR), and mineralocorticoid receptor (MR)) in the different areas of the gland. Data, expressed as the mean ± SD, were evaluated by one-way analysis of variance with Tukey’s post-test (p < 0.05). CR in pregnancy increased the amount of GR, MR, and Ki-67 receptors in the adrenal gland. The npRC group showed highest GR staining compared to the animals that received a normal diet. Protein restriction in pregnancy decreases adrenal MR. The results allowed us to conclude that even without altering the weight of the adrenal glands, the pRC group suffered the most from stress during the study, suggesting that CR associated with pregnancy can cause morphofunctional changes in the adrenal glands.

## 1 INTRODUCTION

Adrenal glands (AGs) are bilateral structures located above the upper pole of the kidneys. In healthy young women, the GA is pink, smooth, and opaque. In men, it is a smaller and more reddish organ that appears slightly translucent. When sectioned, the tissue is soft, and the pith, with a rounded to oval appearance, is centrally placed and dark red in color. In terms of weight, the adrenal gland is approximately 25% heavier in women than in men, as women have a wider cortex, but the medulla is the same size in both sexes (Hardy & Cooper, 2010). Histologically, it is divided into the cortex and medulla and acts as an essential regulator of the stress response (Kanczkowski, Sue & Bornstein, 2017). In rats, the adrenal glands are paired above the kidneys; the right adrenal gland is somewhat medial to the superior pole and intimately connected to it, while the left GA is above this organ (Dunn, 1970).

Calorie restriction (CR) is defined as a reduction in caloric intake below usual *ad libitum* without malnutrition, which generally represents a 10–40% decrease in caloric intake with no reduction in the nutritional content of the diet (Bagherniya *et al*., 2018). This results in a delay in aging, prolongation of the maximum and average lifespan in animals of different species, and a significant decrease in cardiovascular diseases, diabetes, neurodegenerative diseases, and cancers (Al-Regaiey, 2016).

Protein restriction (PR) can be defined as restriction of amino acid intake without malnutrition (Youngman, 1993). Studies on yeast and flies have shown that amino acid restriction promotes longevity and protection. In rodents, protein restriction prolongs lifespan and alleviates harmful phenotypes associated with aging (Mirzaei; Raynes & Longo, 2016).

Other significant health benefits of nutritional restriction have also been demonstrated, including decreased tumor angiogenesis (Hawrylewicz et al., 1982; Youngman & Campbell, 1992), antioxidant enzymes that enhance defenses (Lammi-Keefe et al., 1984), improved immunologic responses (Jose & Good, 1973; Bell et al., 1990), and reduction of total serum cholesterol (Terpstra et al., 1981; Youngman, 1987). PR animals generally have a smaller body size (Youngman & Campbell, 1992) and more physically active (Krieger et al., 1988). Furthermore, both PR (Youngman; Park & Ames, 1992) and RC (Lok et al., 1990) significantly decrease cell division rates in many tissues (Youngman, 1993).

During the gestational period, numerous changes occur in the body to meet the needs of the mother and fetus. Among these changes are the accumulation of maternal adipose tissue, increased metabolism rate along with increased cardiac output and respiratory rate, and a higher calorie intake, which is essential for having a pregnancy that does not bring risks to the pregnant woman, let alone to the offspring (King, 2000).

During pregnancy, nutrient metabolism undergoes adjustments caused by hormonal changes, the demand of the fetus, and maternal supply of nutrients, especially during the last half of pregnancy, which is the period in which the fetus grows the most. These transitions, along with behavioral habits, changes in the amount of food consumed or energy expended, food choices, or type of physical activities of mothers, increase the physiological adjustments necessary during pregnancy. However, when the physiological adaptation of the body is exceeded during this phase, fetal development can be harmed (King, 2000).

Deficient nutrition during pregnancy results in permanent metabolic and/or structural adaptations in the offspring. Females with caloric deficiencies or malnutrition during pregnancy affect their offspring, increasing the risk of developing pathologies in adult life, such as metabolic syndrome, obesity, cardiovascular diseases, and type 2 diabetes mellitus (Chango & Pogribny, 2015). It is known that pregnant females can suffer from increased blood pressure (Gao; Yallampalli & Yallampalli, 2012), along with changes in the immune system (Thiele; Diao & Ark, 2017), as well as being subject to changes in their AG during the gestational period.

Although progressive, there has been a reduction in the number of malnourished individuals in today’s society. The dietary patterns of the contemporary society have undergone changes due to advances in food production and industrialization technologies. Other factors have also changed, such as the assimilation of cultural patterns and modification of life habits. Thus, today, important issues related to the impact of nutritional restriction on metabolism should be evaluated in animal models with dietary restrictions. The changes in GA during pregnancy in rats are poorly understood. Given the importance of this gland in the production of hormones in females, describing the changes induced by food restriction and pregnancy is essential to understand its physiology. Therefore, this study aimed to evaluate the morphofunctional organization of the adrenal gland in young adult Wistar rats subjected to nutritional restriction (CR and PR), regardless of the pregnancy status.

## 2 MATERIALS AND METHODS

### 2.1 Experimental procedure

The study was carried out in accordance with the rules established by the Arouca Law, approved by the ethical principles of animal research adopted by COBEA and by the Ethics Committee on Animal Use of the Centro Universitário da Fundação Hermínio Ometto, FHO, opinion 062/2016. Female Wistar rats (10 weeks old, weighing approximately 250–300 g) were subjected to mating. Once the presence of spermatozoa in the vaginal lavage was verified, these animals were called the pregnant group (p, n = 15) and the other group of non-pregnant rats was called the non-pregnant group (np, n = 15). After separating the groups, the rats were divided into three subgroups: those that received a normoproteic control diet (C, 17% casein, n = 5), an isocaloric low-protein diet (PR, 6% casein, n = 5), or caloric restriction of 50% (CR, 50% of the diet consumed by group C, n=5) for a period of 21 days. Diets were calculated daily, considering the weight of the amount offered and the amount that was left for the controls, that is, the amount ingested by the control group. From this, 50% was calculated for the RC group. The rats were kept in individual cages in a temperature-controlled environment (21 ± 1° C) with a 12 h light/dark cycle and free access to water. On the 21st day of gestation (21dG, P animals) or on the 21st day of diet (NP groups), after deep anesthesia with ketamine (100 mg/kg) and xylazine (10 mg/kg), the animals’ right adrenals were collected, weighed, and processed for structural analysis.

### 2.2 Body growth and food consumption

Rats were weighed once a week on days 0, 7, 14, and 21 of the experimental (or gestational) study. The diet was weighed daily for 21 days. In addition to the growth curve, the mass gain after subtracting the initial masses was determined. Food consumption was determined by the difference between the weight of the feed added and the feed remaining in the cages.

### 2.3 Processing for the histomorphometric study of adrenal

After removal, the adrenals were weighed and immersed in a fixative solution containing 10% formaldehyde in Millonig buffer pH 7.4 for 24 h at room temperature. Then, the pieces were washed in buffer and submitted to standard procedures for embedding in paraffin (Paraplast® -Merck). Cross-sections of 5 µm thick pieces were subjected to hematoxylin-eosin staining. Three samples were used for each of the five sections obtained from the median region of each of the three animals in each treatment. Biopsy images were captured on a Leica DM2000 microscope using Leica Application Slite software (version 3.3.0).

From the images, the Image J program (National Institutes of Health, Bethesda, MD, USA) was calibrated to measure the areas of the cortex and medulla, measuring the scale bar divided by its value (50 µm), which resulted in a value of 8.68 µm/pixel. After this process, measurements were started by contouring and measuring the total area and medullary area. The total area was subtracted from the medullary area, which gave the adrenal cortical area. After the measurement, the following ratios were calculated: cortical area/total area, medullary area/total area, and adrenal mass/animal mass. The entire process was performed for all groups, and the results were statistically compared.

### 2.4 Quantification of connective tissue (collagen) in the adrenal by Mallory’s trichrome staining

Cross-sections of 5 µm thick pieces were stained with Mallory’s trichrome stain. Three samples were used for each of the five sections obtained from the median region of each of the three animals in each treatment. Biopsy images were captured on a Leica DM2000 microscope using Leica Application Slite software (version 3.3.0). The images were analyzed using Image J software (National Institutes of Health, Bethesda, MD, USA) by color deconvolution and statistically analyzed.

### 2.5 Processing for immunohistochemistry analyses

We evaluated the expression of glucocorticoid receptors (GR), mineralocorticoid receptors (MR), and Ki-67 antigen in the adrenal glands of pregnant and non-pregnant young adult rats subjected to different nutritional protocols. All procedures were performed according to the protocol established by Gianchini *et al*. (2007). Briefly, antigen retrieval was performed by immersing the silanized slide in sodium citrate solution (10 mM, pH 6.0) for 40 min at 95°C. Each step was followed by washing with PBS. All steps were performed in a humid chamber under care to avoid dehydration of the sections. Incubation with the primary antibody was performed by incubating the sections with anti-GR (monoclonal mouse, Santa Cruz, USA), anti-MCR (monoclonal mouse, Santa Cruz, USA), and anti-Ki-67 (monoclonal mouse, Santa Cruz, USA) antibodies which were diluted 1:200 in PBS containing 3% bovine albumin (v/v) overnight at 4°C. After the primary antibody reaction, a Novolink Polymer Detection Systems Kit (RE7280K; Leica Biosystems Newcastle L10, Newcastle Upon Tyne, UK) containing the secondary antibody was used. After washing in PBS, the peroxidase reaction was visualized using DAB (3,3’-diaminobenzidine) from the same kit. For each immunohistochemical reaction, a negative control of the adrenal sections was performed, omitting the primary antibody. The sections were examined using a Leica DM2000 Photomicroscope in images digitized with the support of the Sigma Scan Pro 5.0™ program and evaluated by area (µm^2^) using the Image J software (National Institutes of Health, Bethesda, MD, USA). Quantification was based on the decomposition of the immunohistochemical image into three base colors: brown (immunohistochemistry), purple (Harris hematoxylin), and green (background of glass slides). Morphometric analysis, corresponding to brown color, was performed using the threshold function (ImageJ), and antibodies/markers were measured as the percentage of total pixels in each image (Landini, Martinelli & Piccinini, 2021). Data are reported as the percentage area of the respective antibody.

### 2.6 Statistical analysis

Data were compared using analysis of variance (ANOVA) followed by Tukey’s post-hoc test using GraphPad Prism software (GraphPad Software, Inc. La Jolla, CA, USA) with a significance level of 5% (p < 0). .05, n = 5). The results were expressed as mean ± standard deviation (X ± SD) and later represented as a percentage of variation in relation to controls, to which a value of 100% was assigned.

## 3 RESULTS

### 3.1 Effect of nutritional restriction on female characteristics

The body mass gain of rats was analyzed by weighing the animals weekly at time 0 (mating day) and on the 7th, 14th, and 21st gestational days (Figure 1). Throughout the study period, the pC group of animals presented an ascending weight curve, indicating that the expected growth occurred during pregnancy, while in the pRC group, there was weight loss during the first 14 days when compared to the initial weight, with mass gain only in the last week of pregnancy. The pRP group also showed a lower body mass gain. There was a gradual decrease in weekly consumption of diet in the groups when evaluated from the 1st to 3rd gestational weeks. Pregnancy and/or diet did not alter the mass in the right adrenal gland.

**FIGURE 1.**
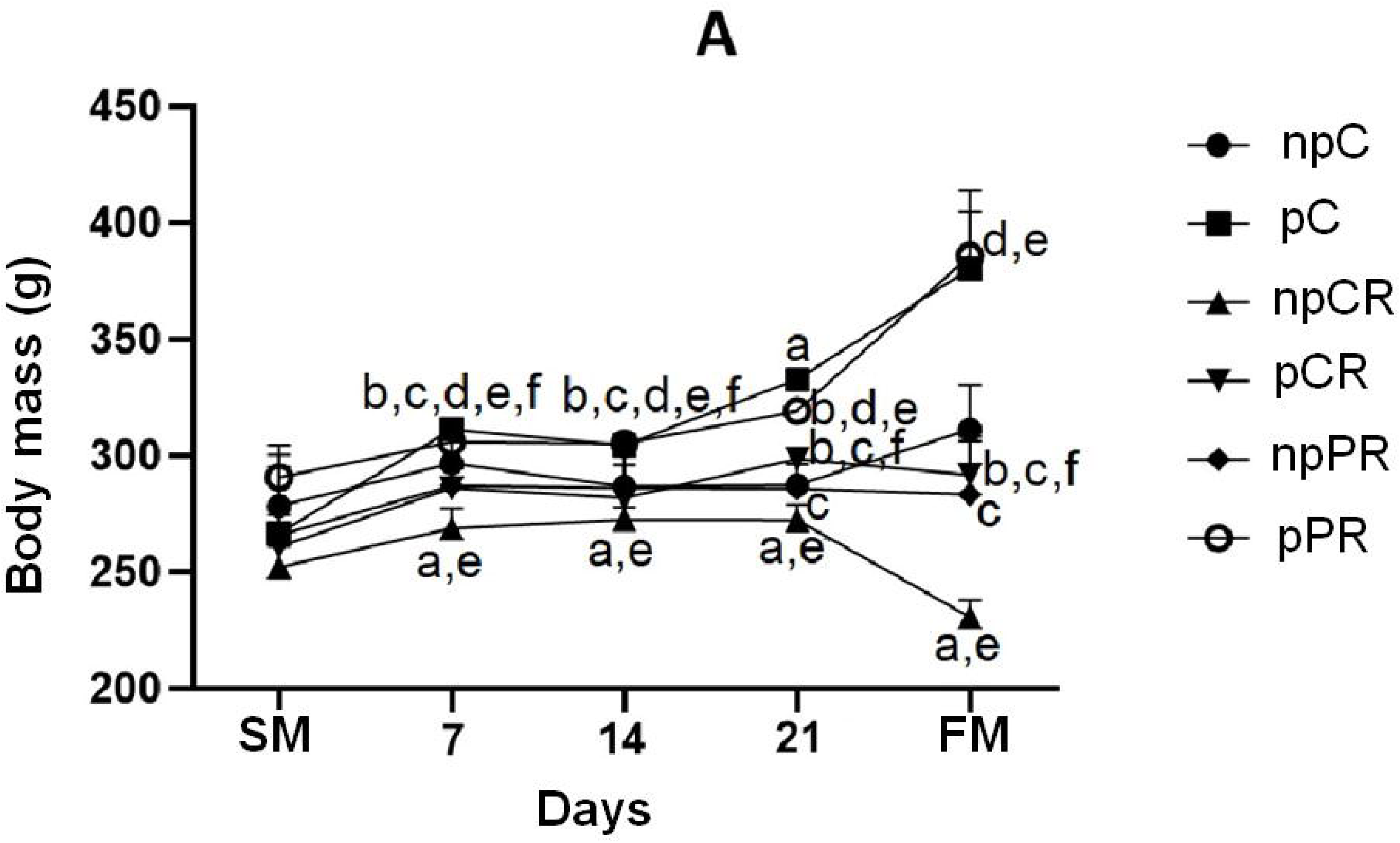
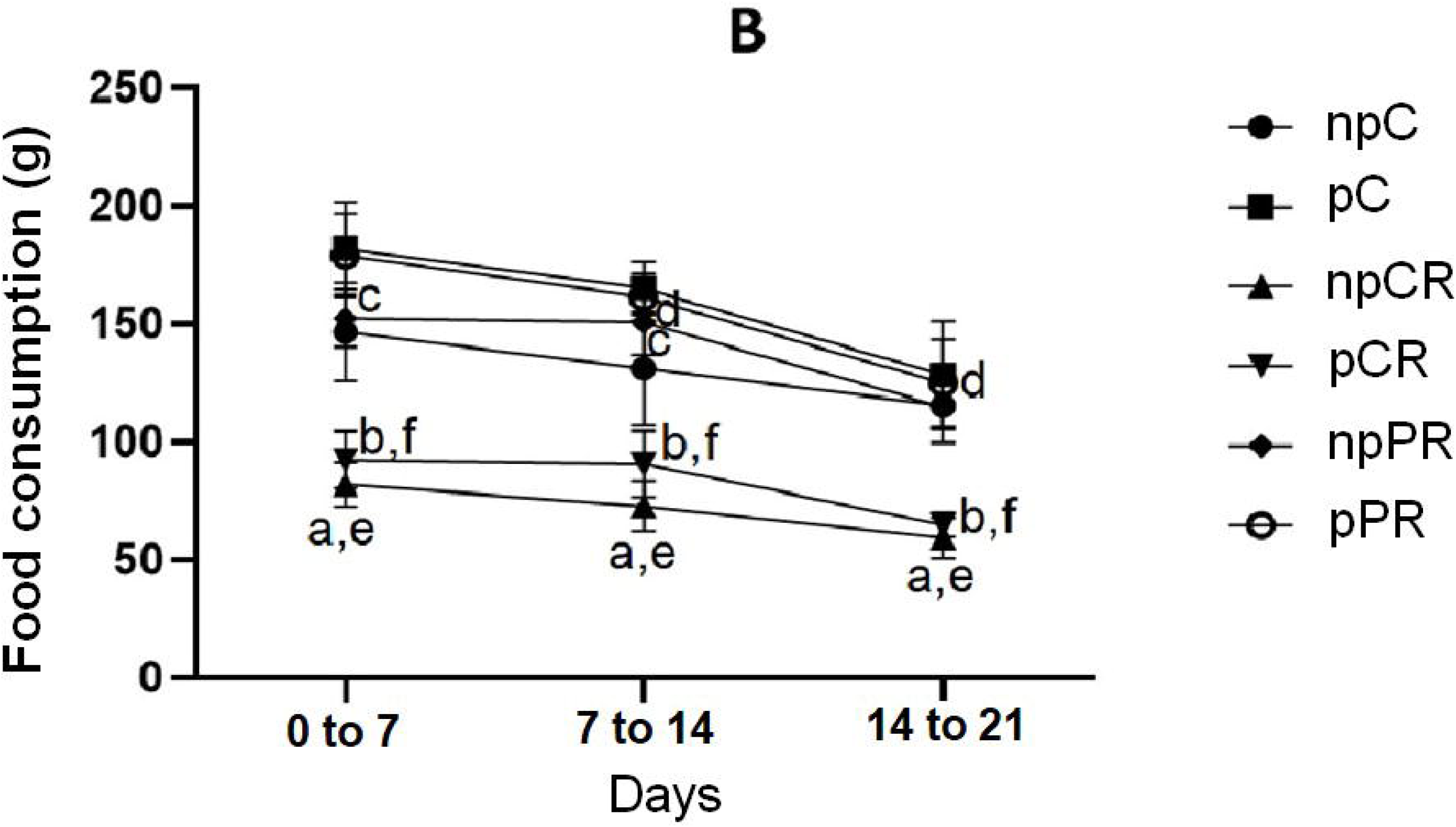
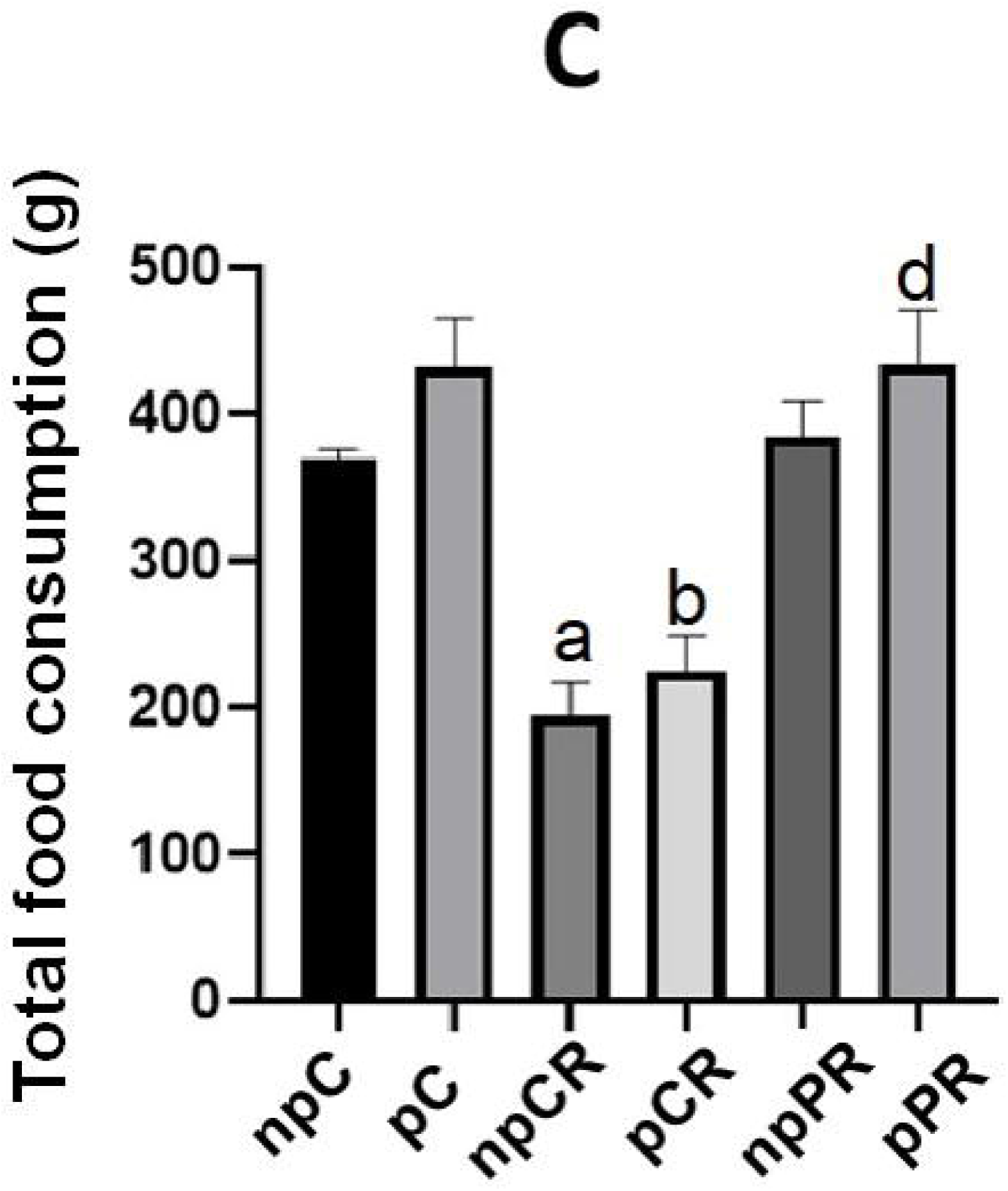
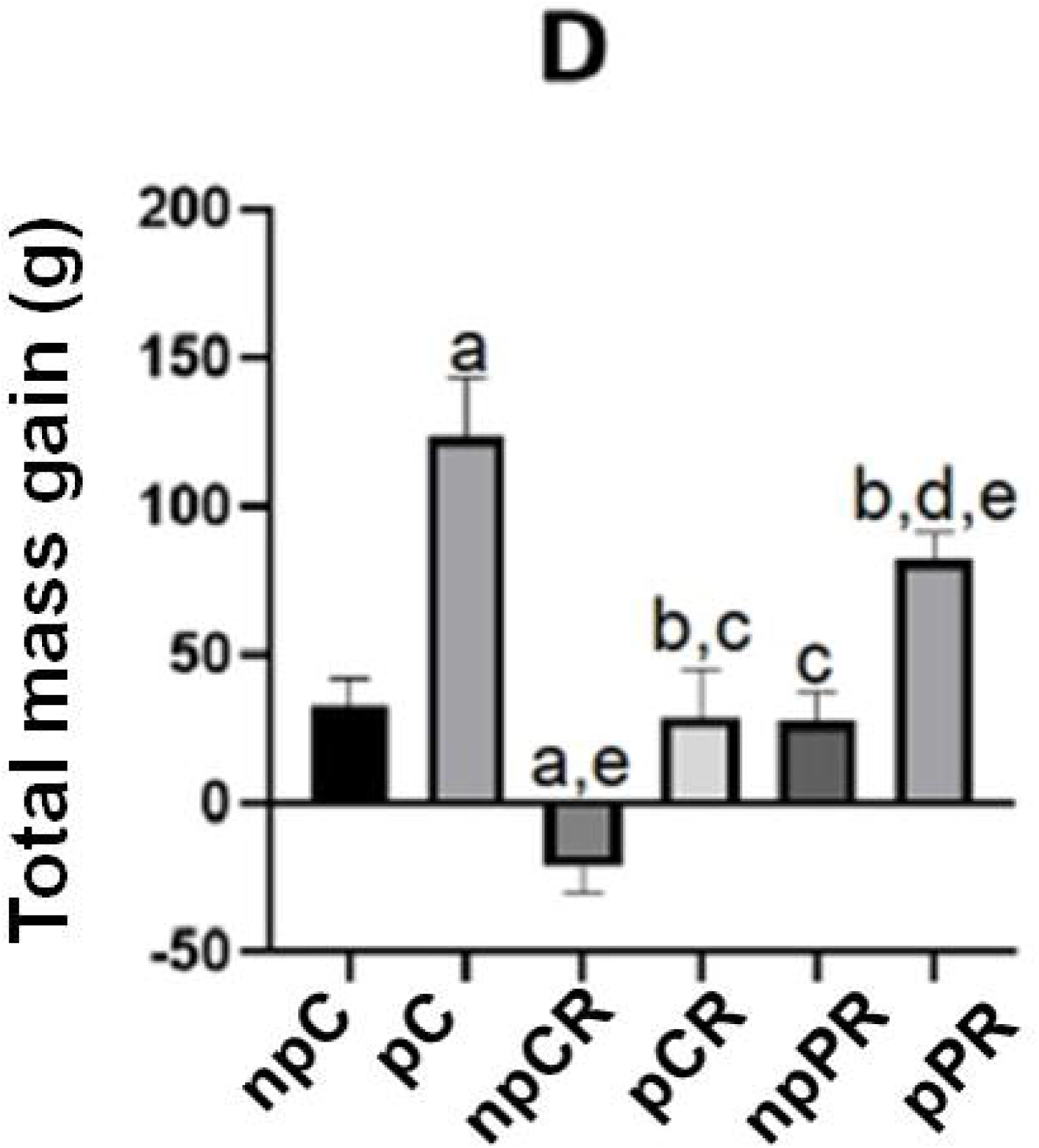
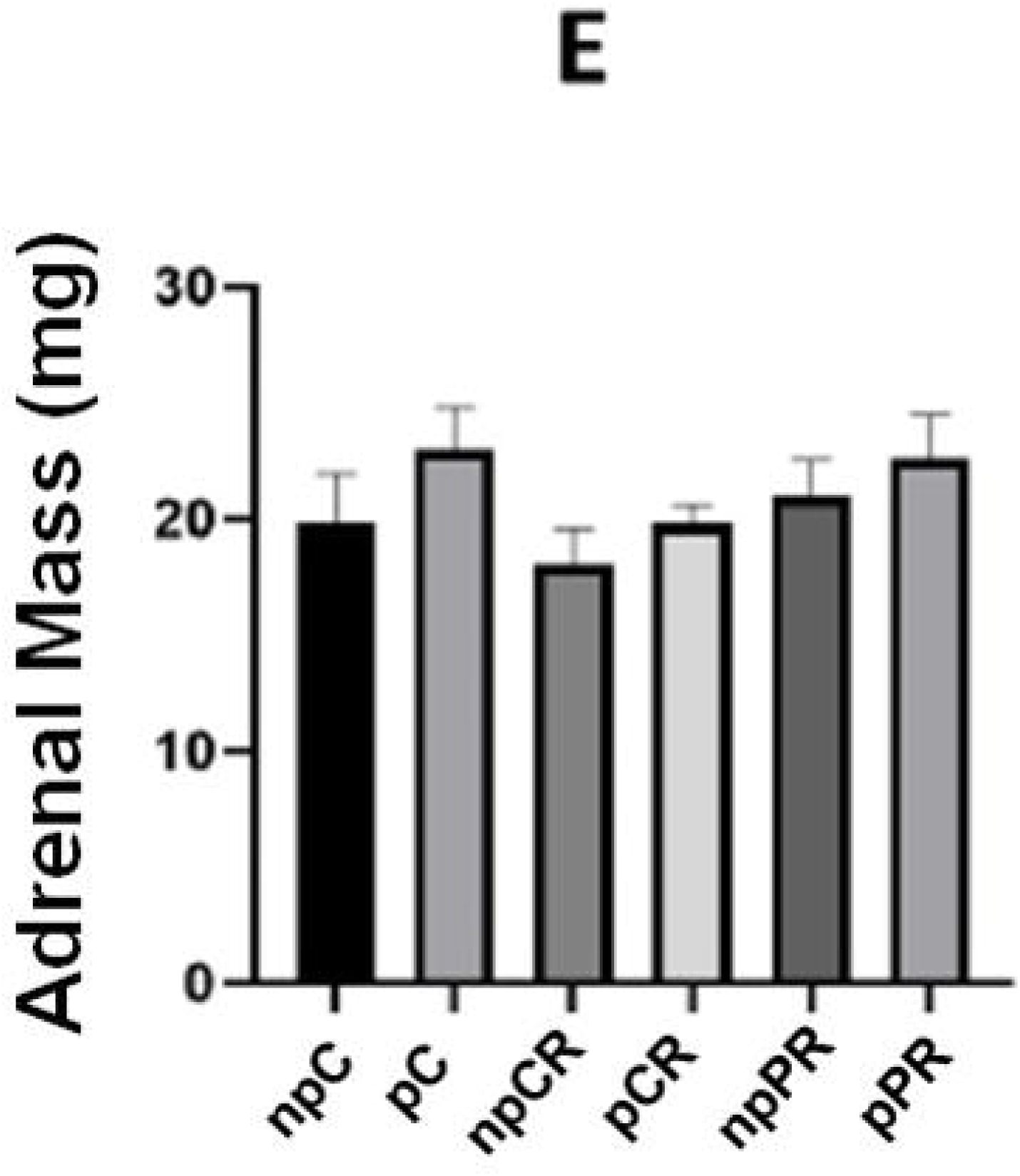
Nutritional restriction in pregnant and non-pregnant rats. A) Body mass of pregnant and non-pregnant animals. B) Food consumption. C) Total food consumption of the animals. D) Total weight gain of the animals (n = 5). E) Mass of the right adrenal glands after euthanasia. Mean ± SD (n = 6; p < 0.05). (ANOVA, post Tukey test).

### 3.2 Effect of caloric restriction on adrenal gland histomorphometry

There was no significant difference between the experimental groups for the cortical, medullary, and total areas. However, the difference was evident in the adrenal mass/animal mass ratio, with the npC, pC, and pRP groups showing lower values than the npRC group. The pRC and npRP groups showed values higher than pRP and pC, which were statistically lower than those of npRP (Figure 2).

**FIGURE 2.**
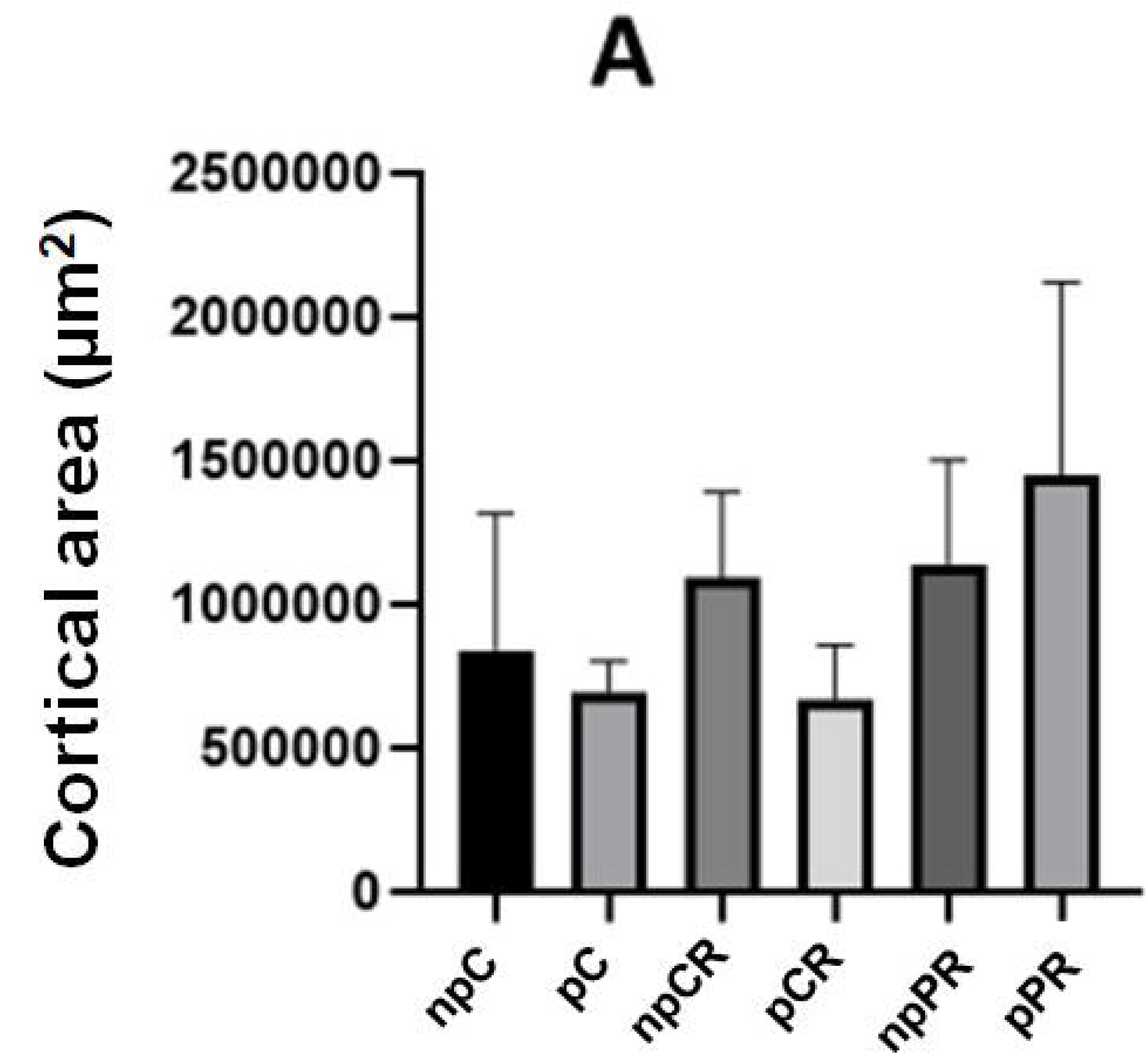
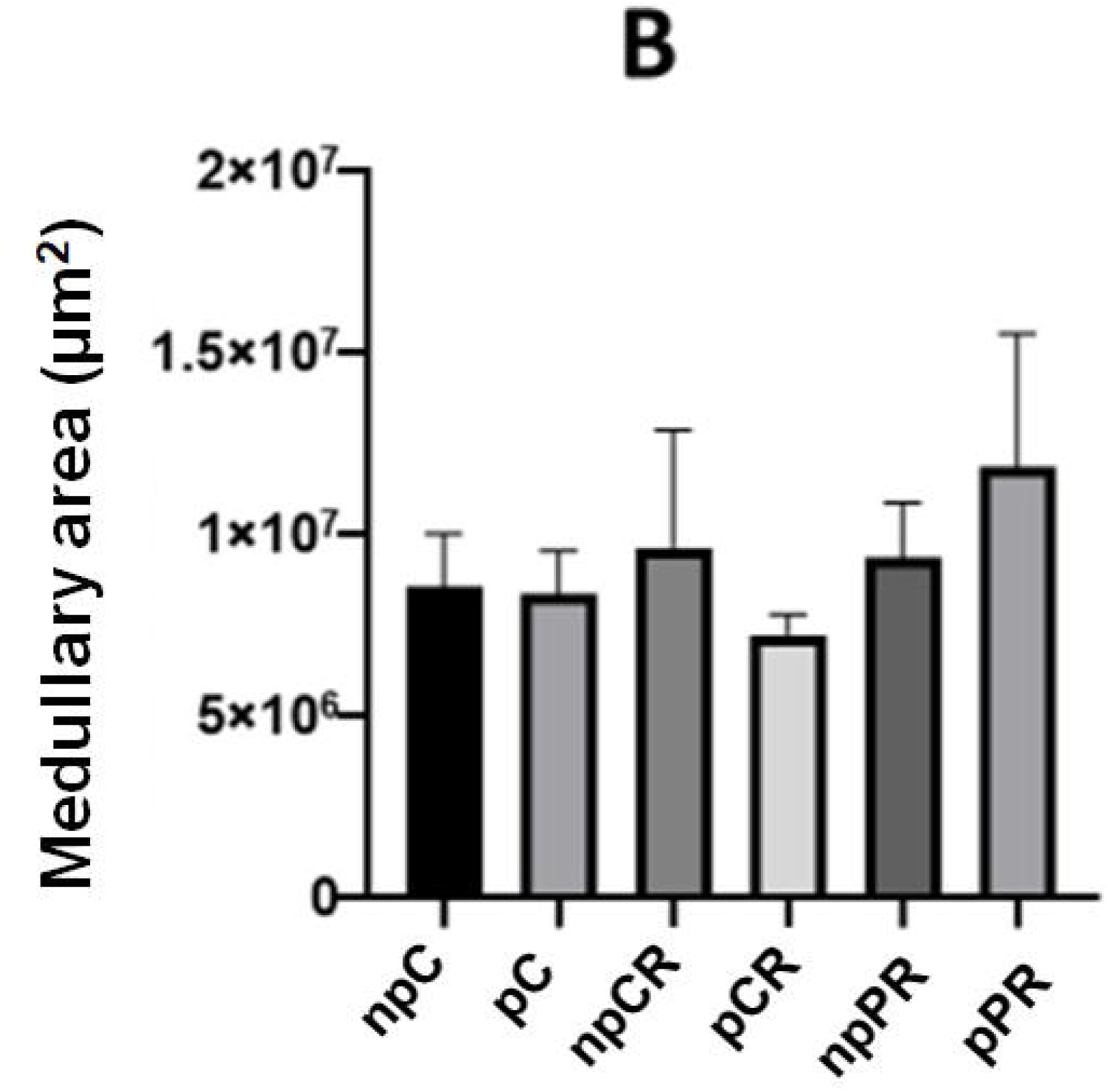
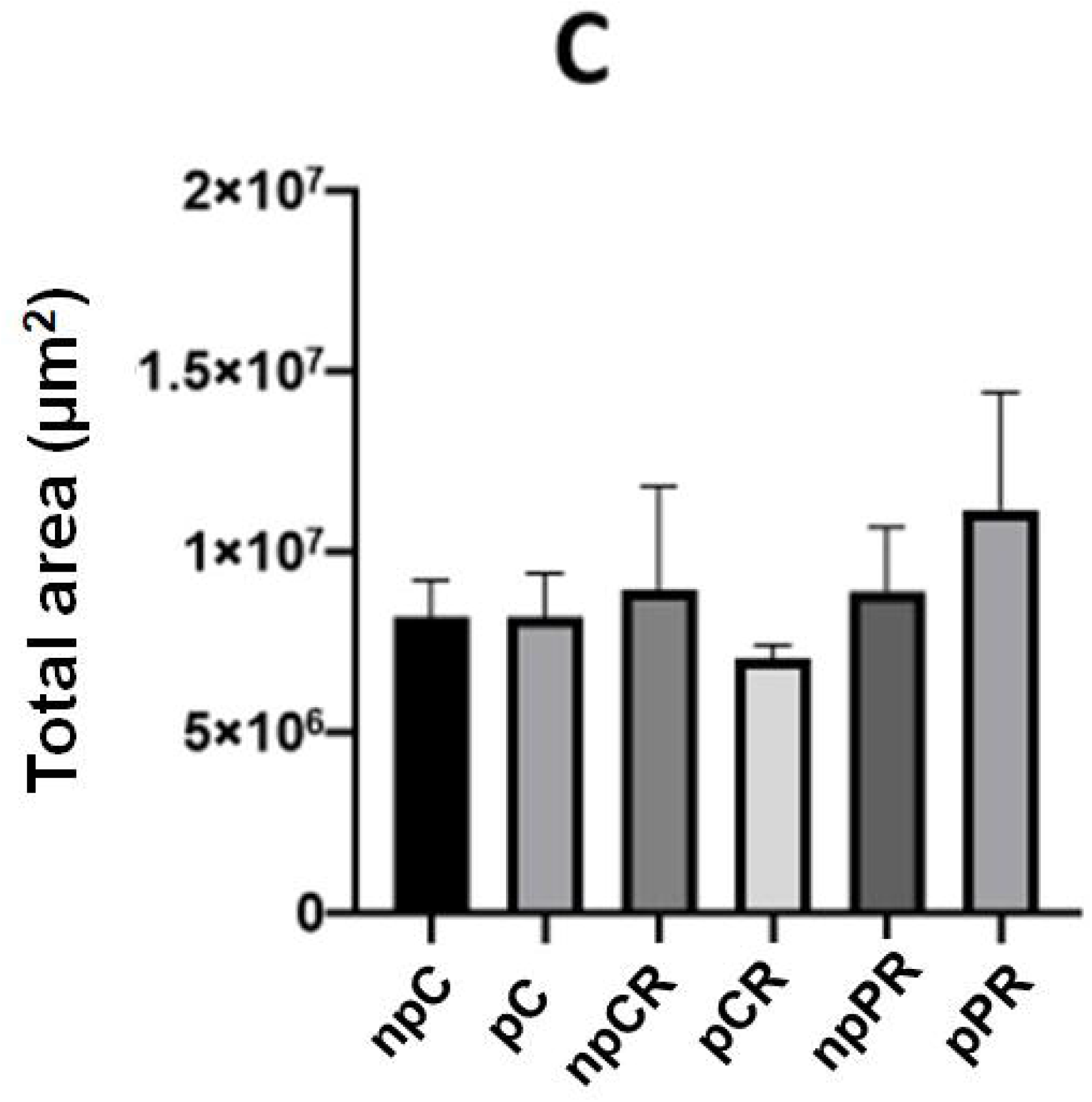
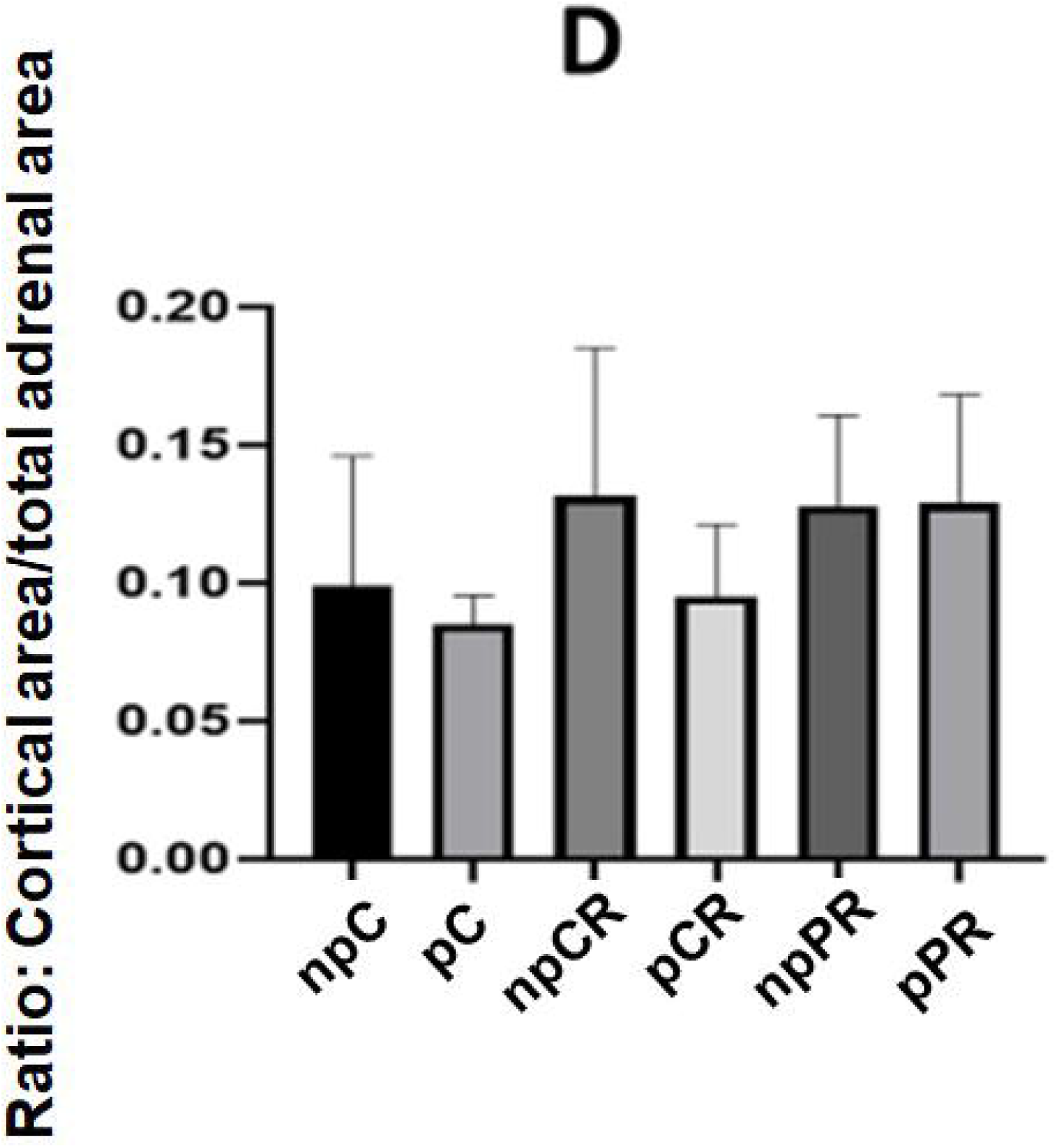
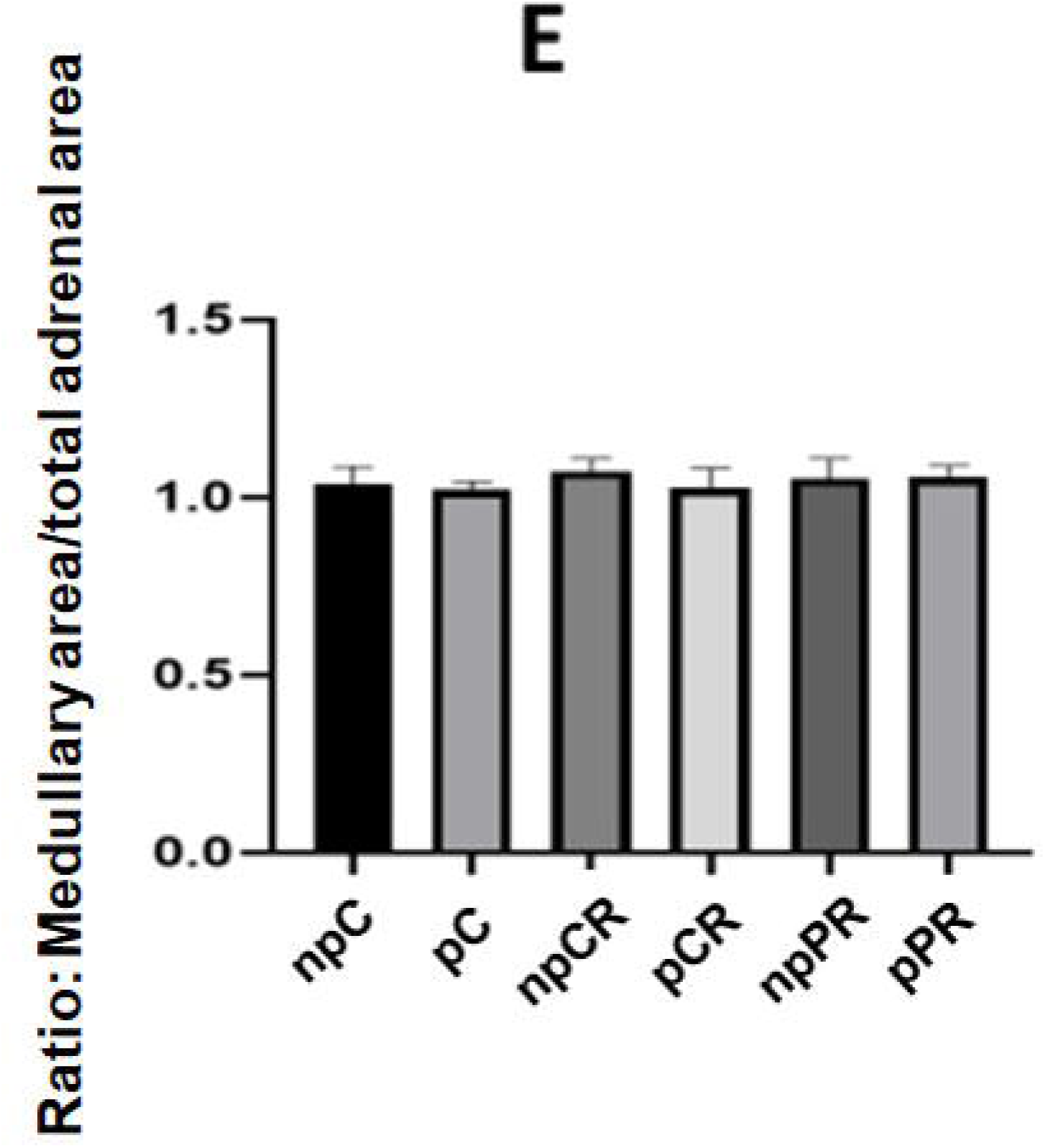
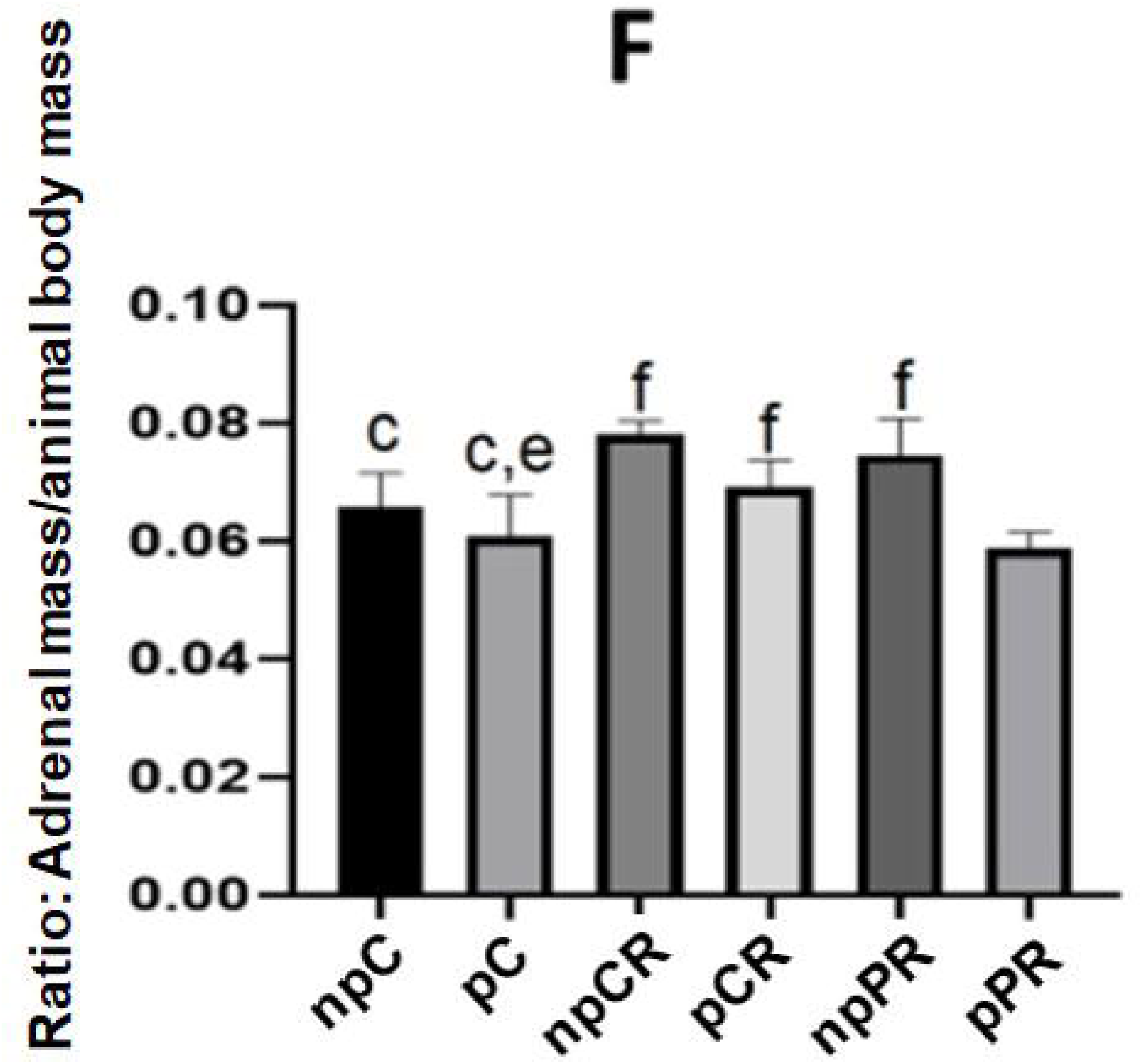
Histomorphometry of the adrenal glands. A) Cortical area. B) Medullary area. C) Total area. D) Ratio of cortical area to total adrenal area. E) Medullar area to total adrenal area ratio. F) Ratio of adrenal mass to animal body mass (n = 5). Mean ± SD (n = 6; p < 0.05). (ANOVA, post Tukey test).

### 3.3 Quantification of Connective Tissue (Collagen) by Mallory’s Trichrome

In the quantification of connective tissue (collagen) by Mallory’s trichrome, the zona glomerularis of the pRC group showed the smallest amount of connective tissue (collagen) area compared to the npC group (Figure 3A). In other areas, there were no differences between the groups (Figure 3).

**FIGURE 3.**
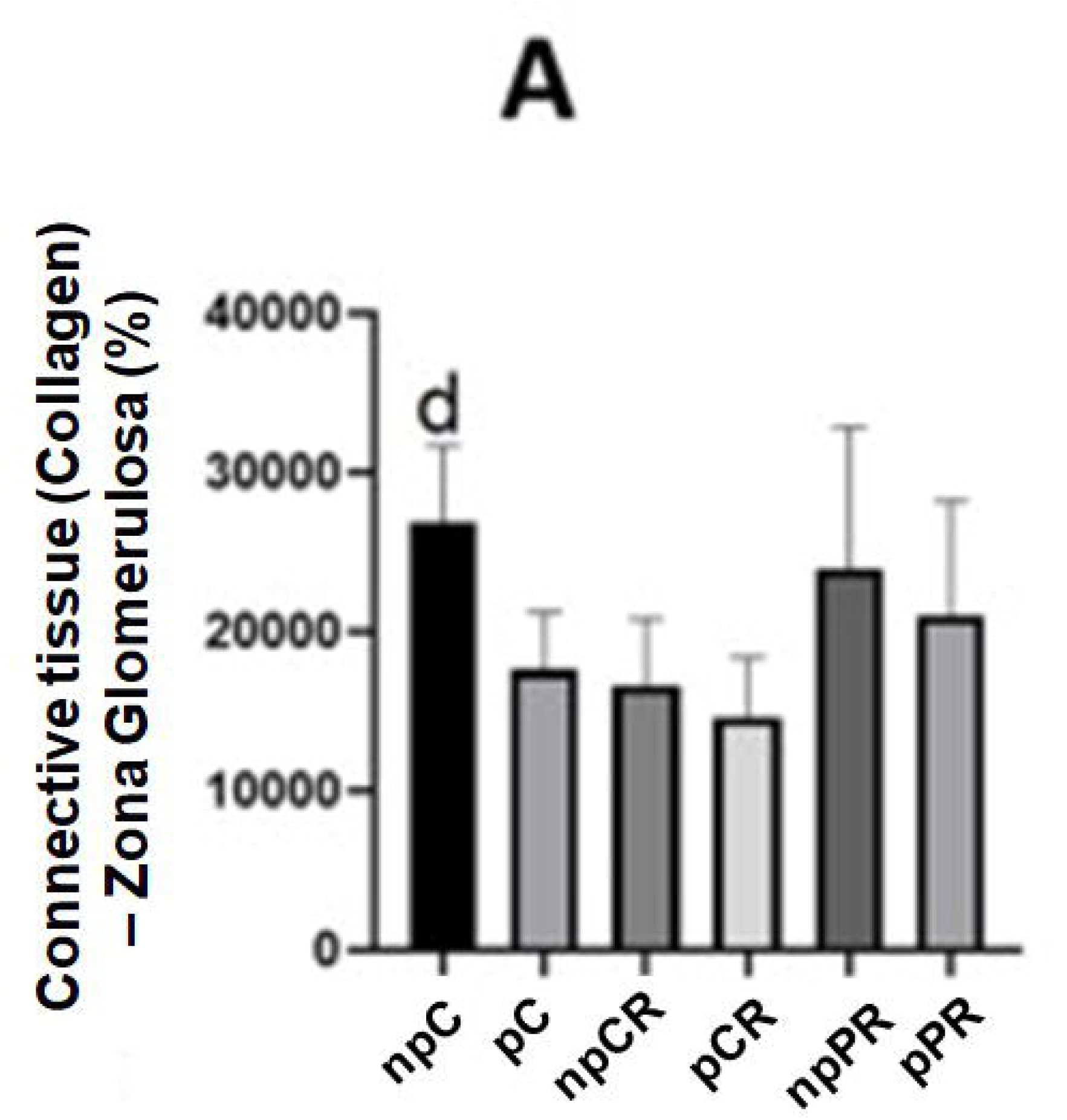
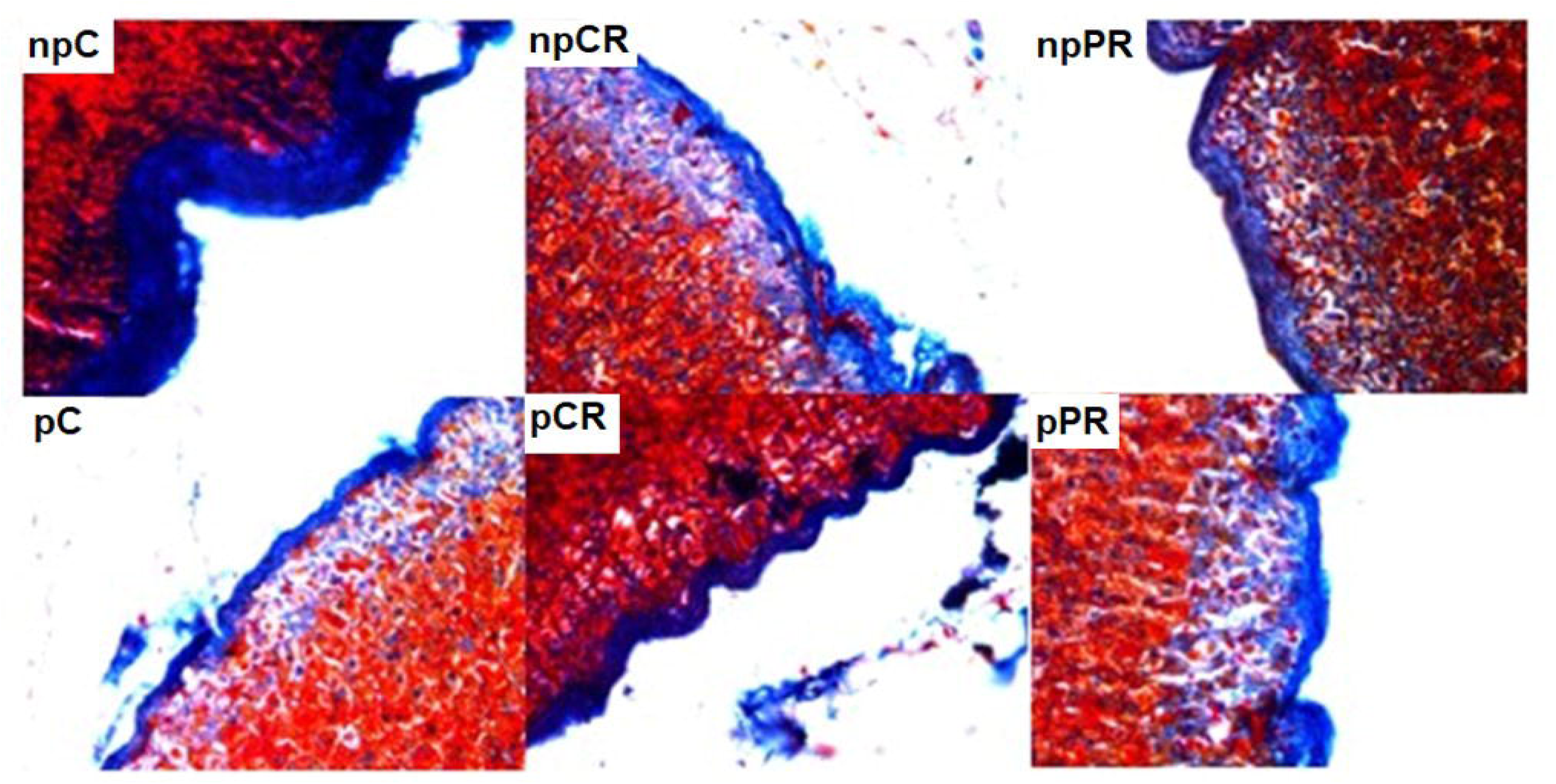
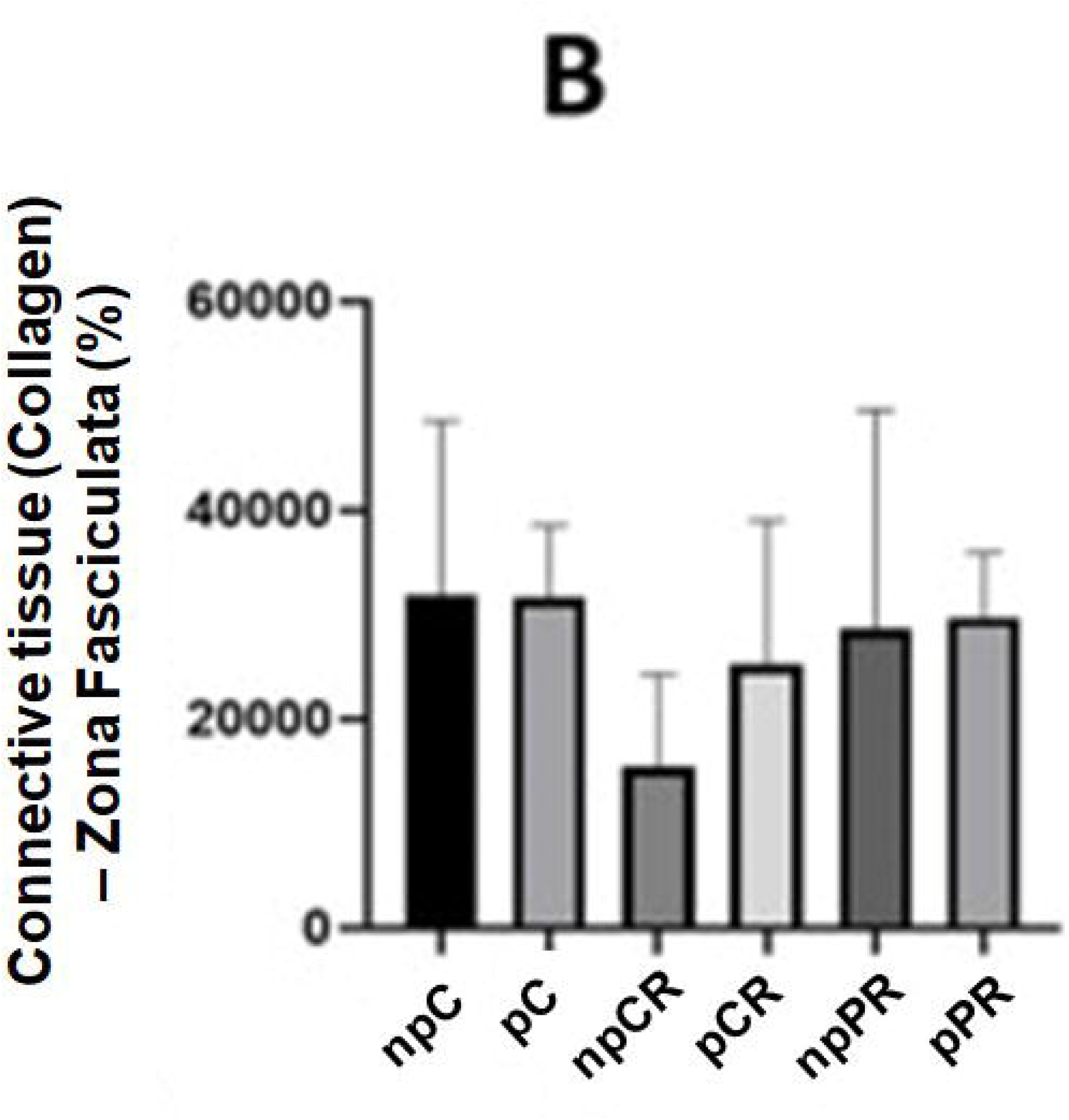
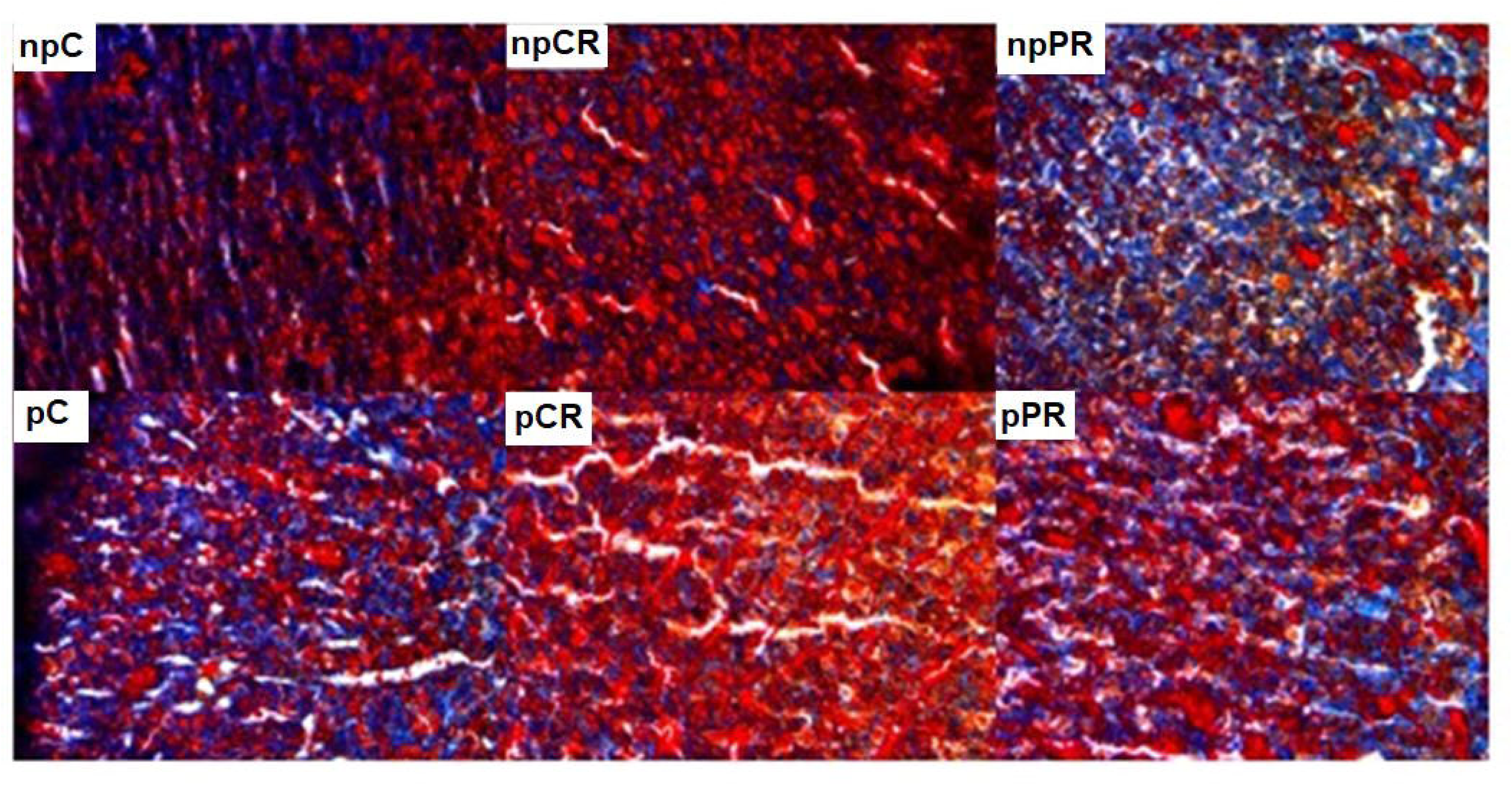
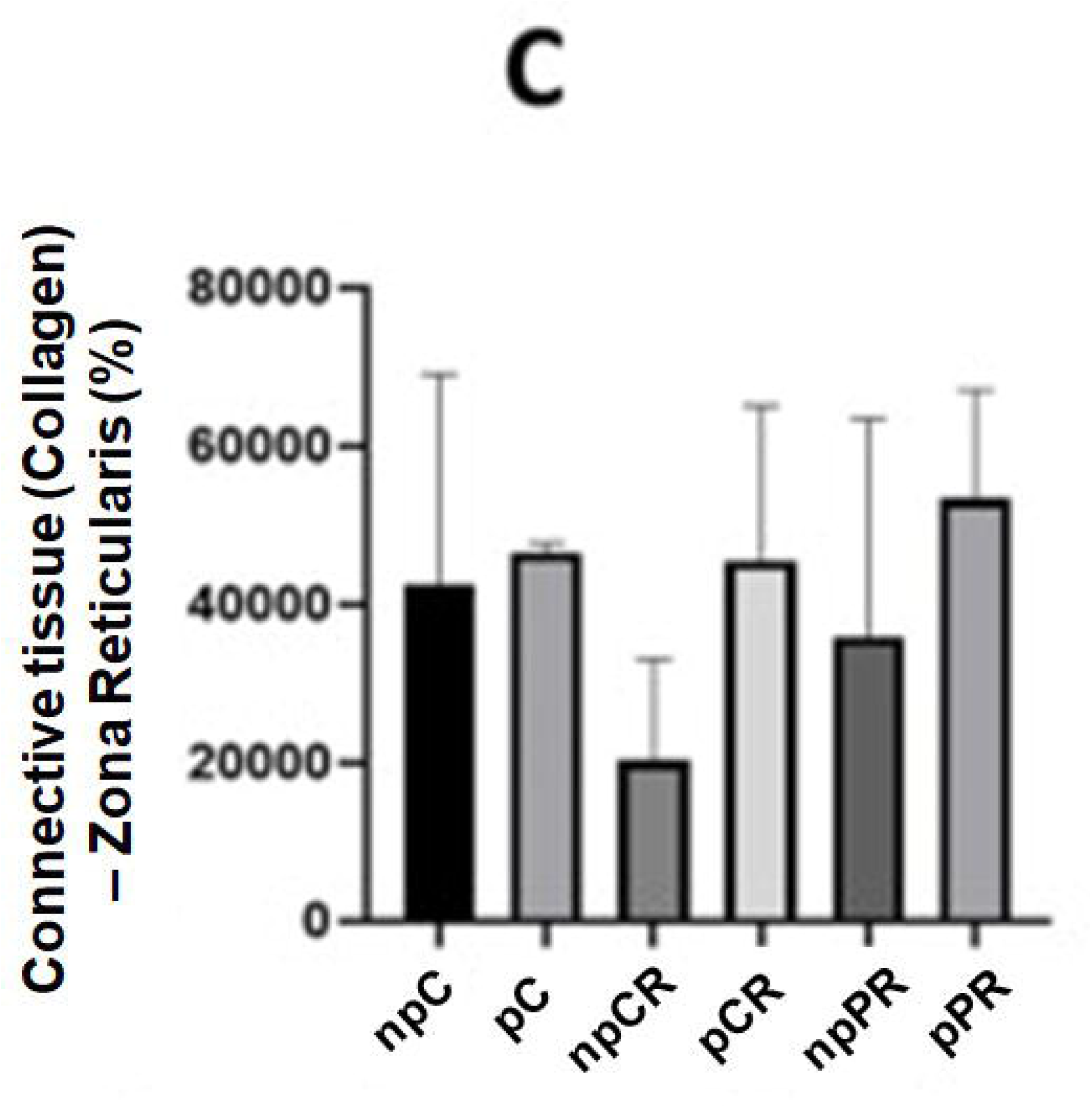
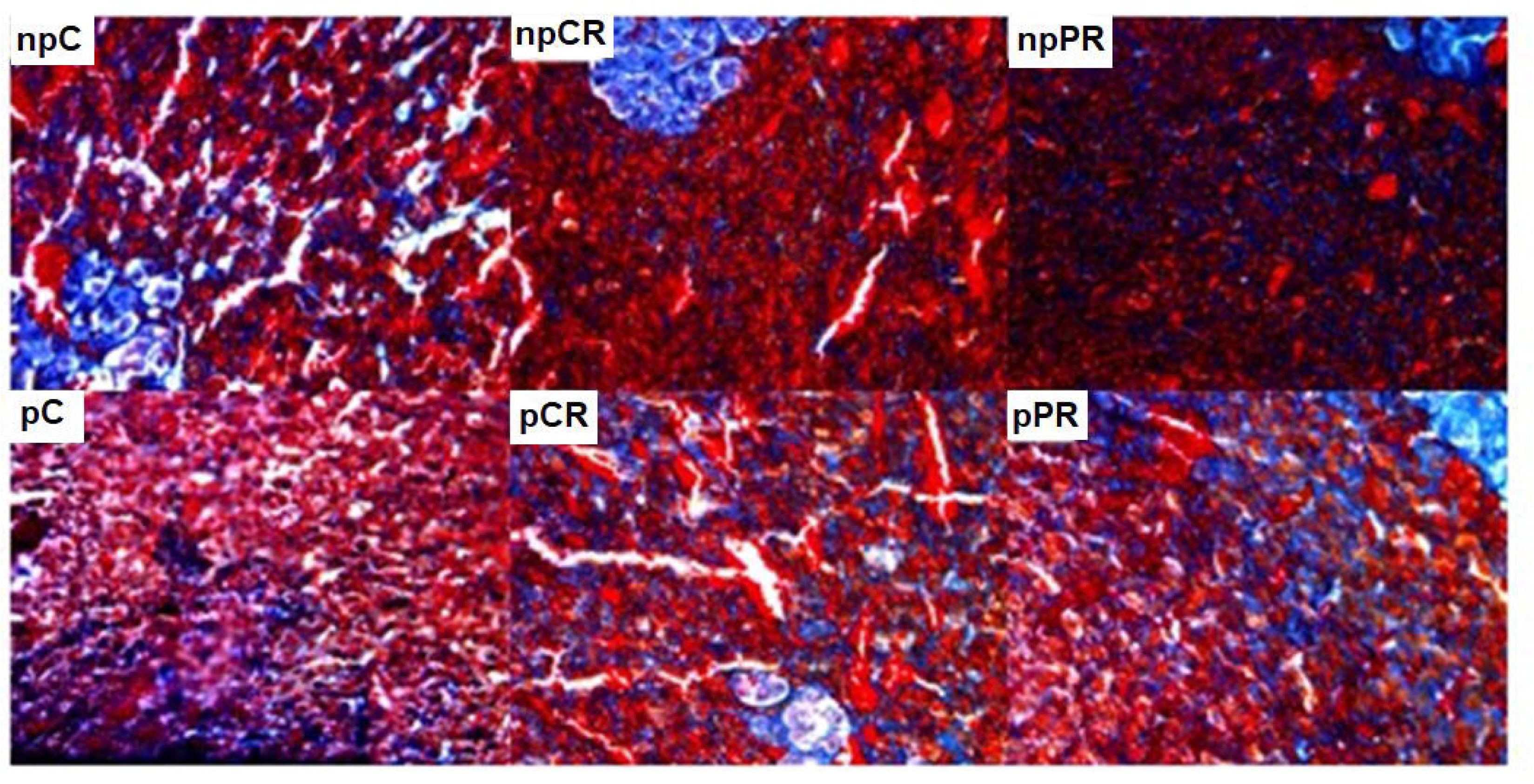
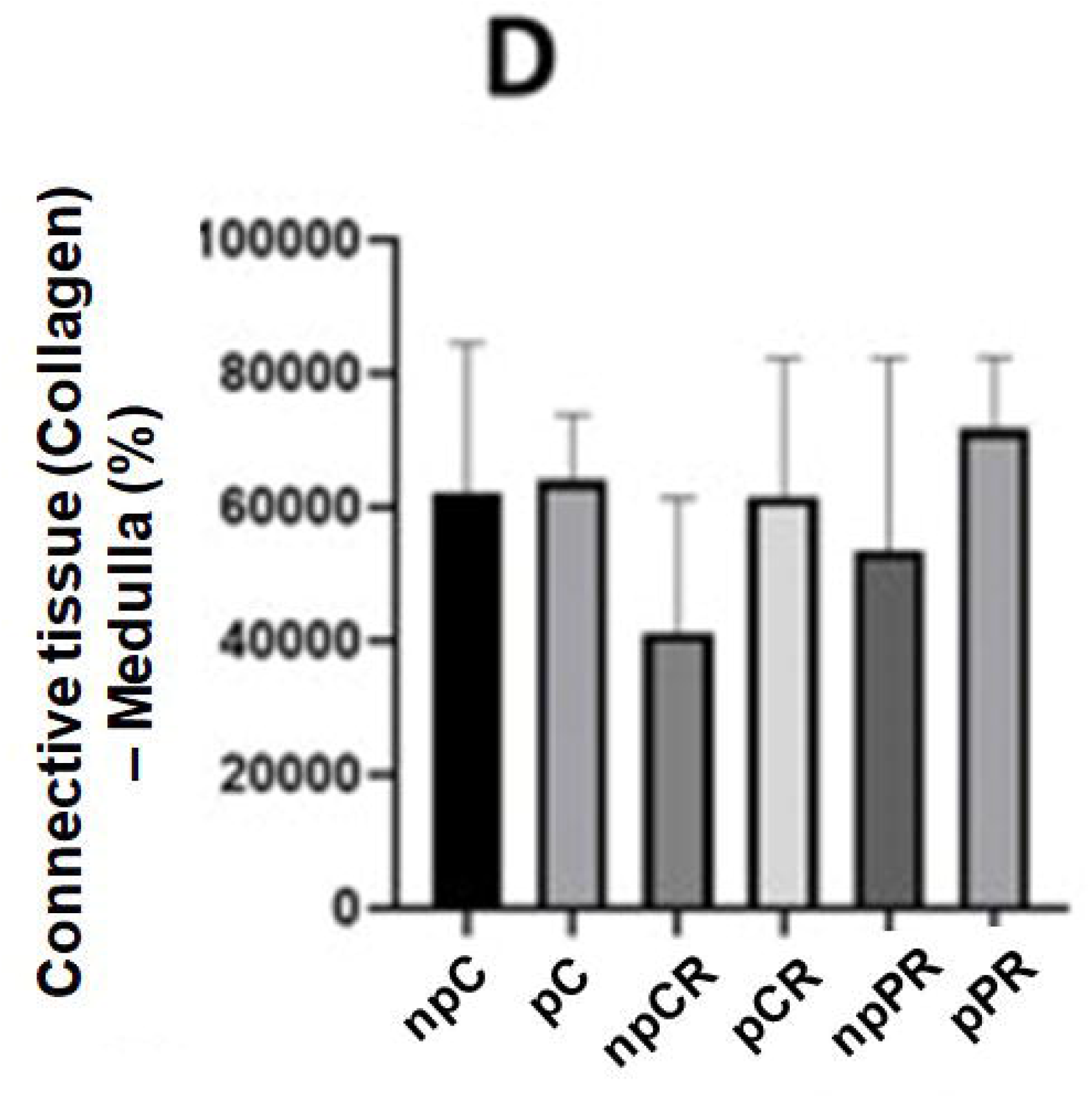
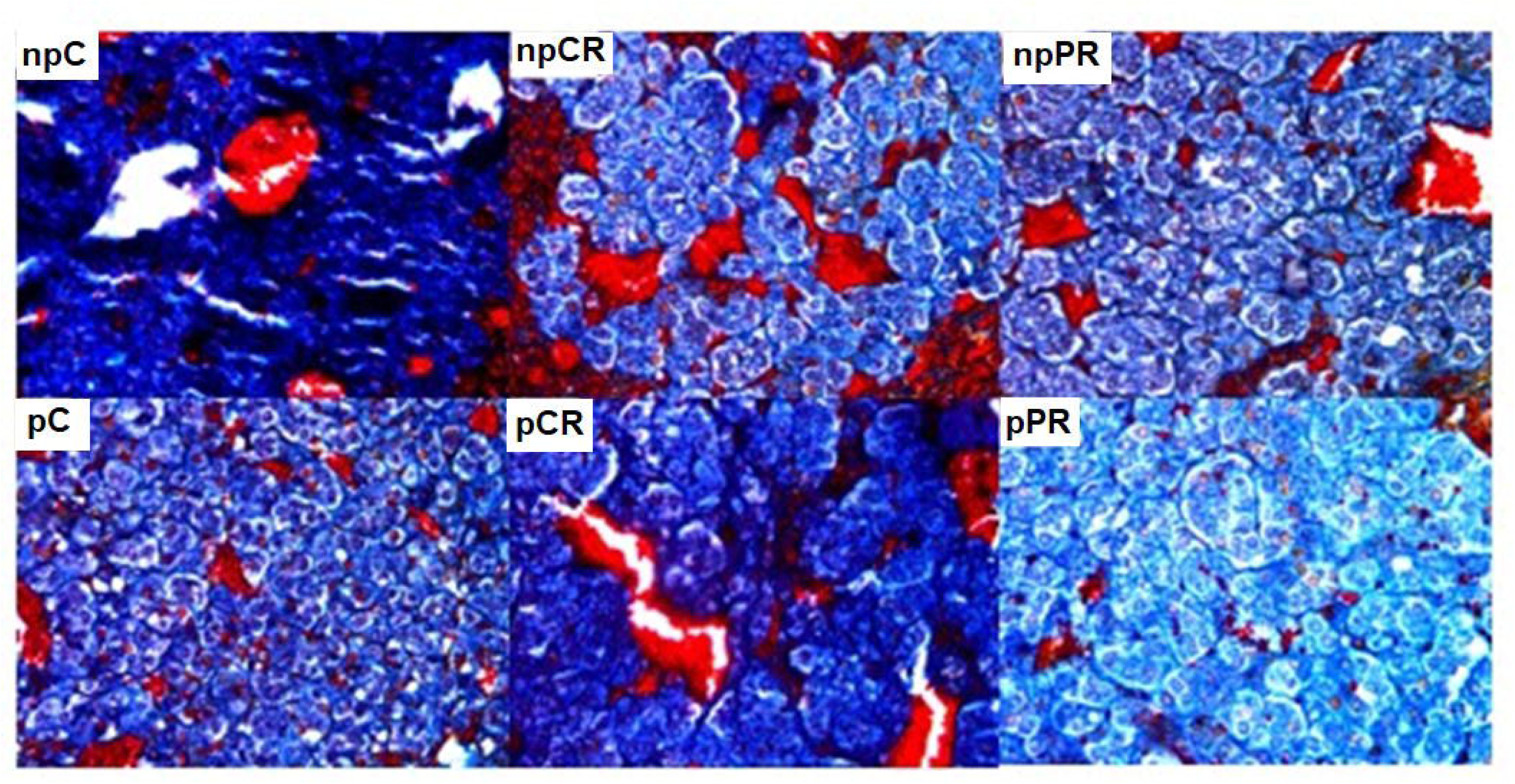
Analyzed area of connective tissue (collagen) in the adrenal gland. Quantification of connective tissue (collagen-in blue), in percentage, in the areas of the adrenal glands using Mallory’s Trichrome staining. Final magnification – 200x. A) Glomerular zone. B) Fasciculate zone. C) Reticulated zone. D) Medullary zone. Mean ± SD (n = 5; p < 0.05). (ANOVA, post Tukey test).

### 3.4 Cell division in the adrenal glands assessed using the Ki-67 marker

The pRP group had a reduced number of dividing cells in the zona glomerularis compared with the pC and pRC groups. Among the control groups, pC showed greater cell proliferation than npC. In the zona reticularis, the nutritionally restricted groups, npRC and npRP, showed higher Ki-67 staining than the npC group. In the reticular and medullary zones, the npRP group had a greater number of dividing cells than the pRP group (Figure 4).

**FIGURE 4.**
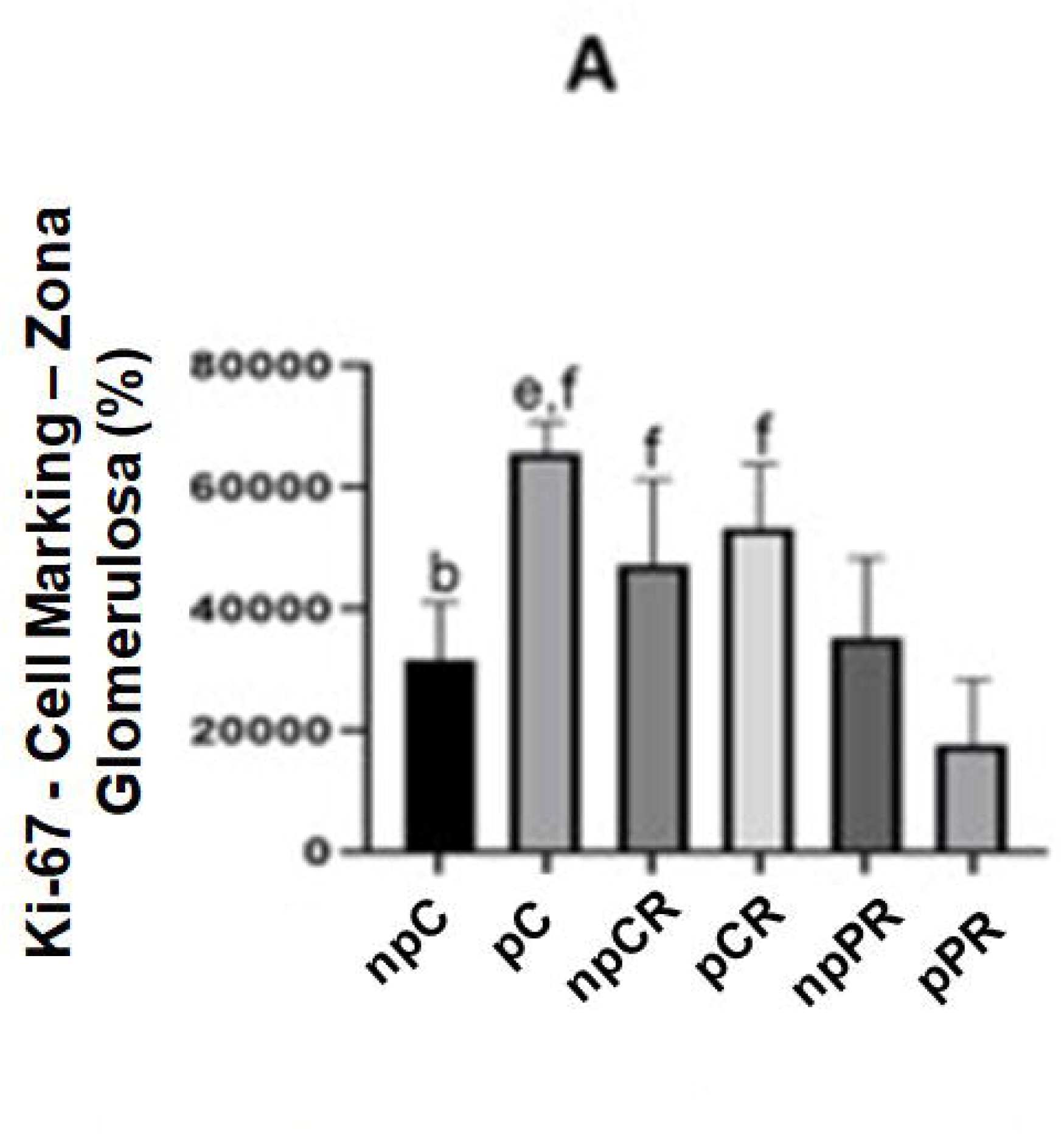
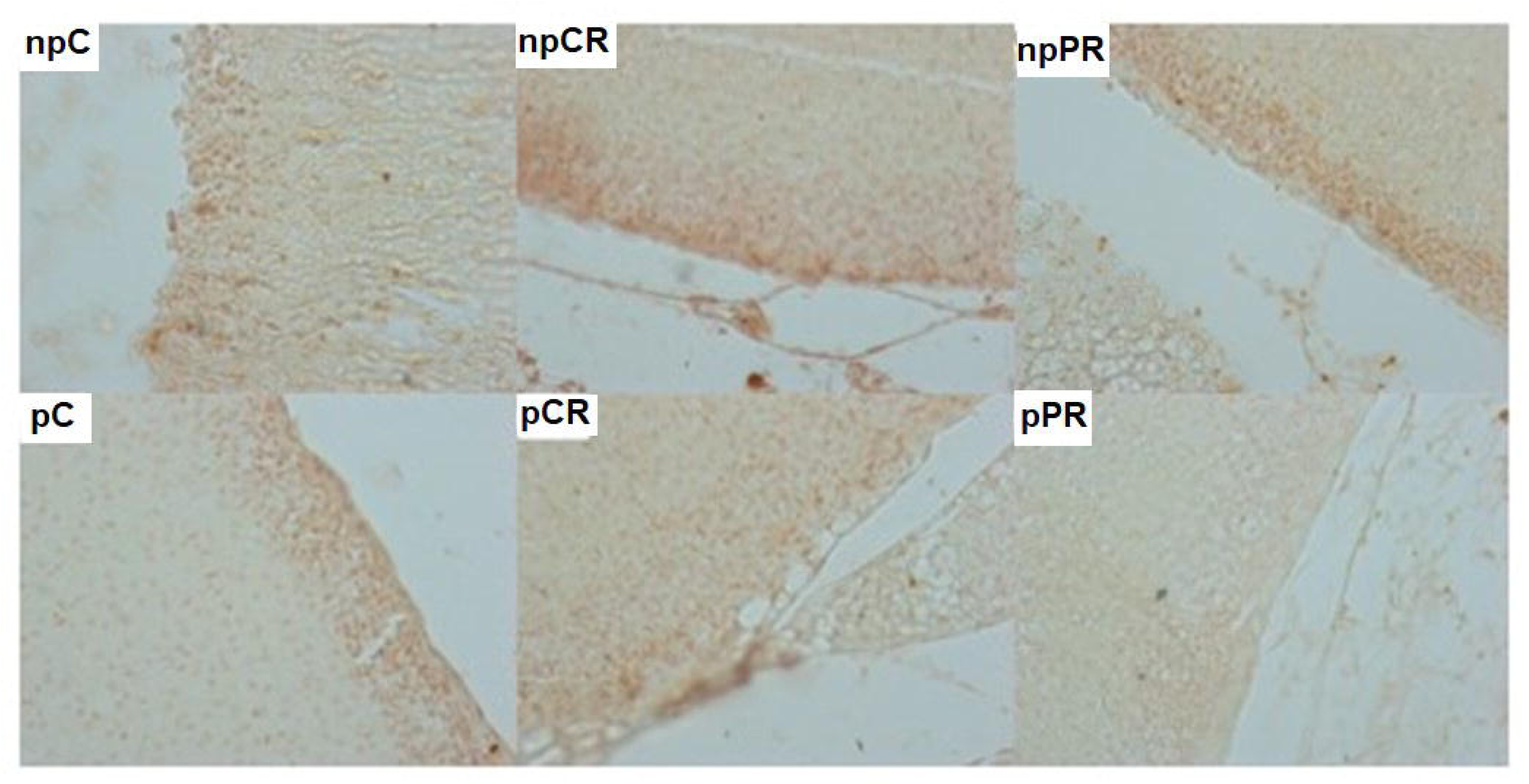
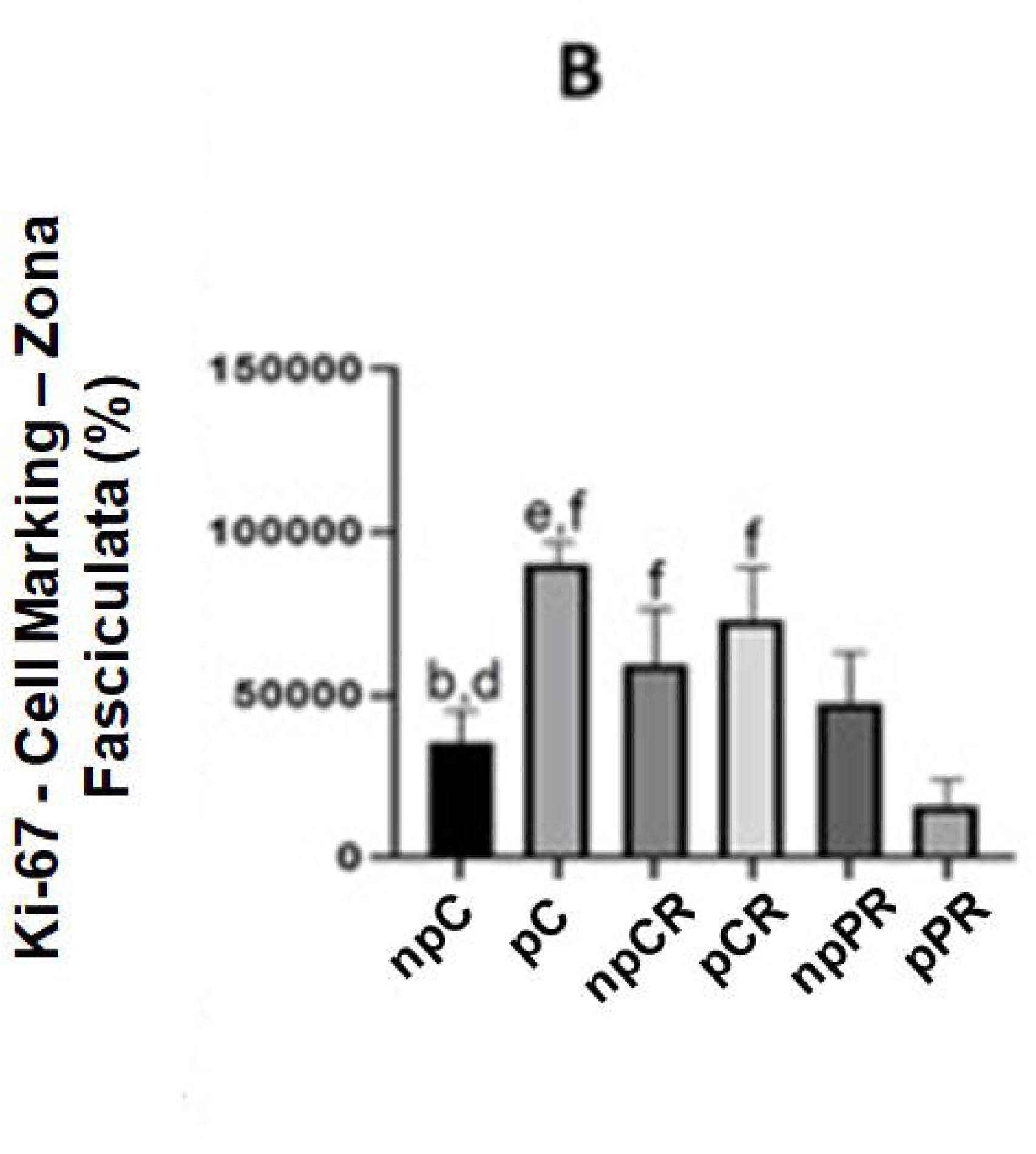
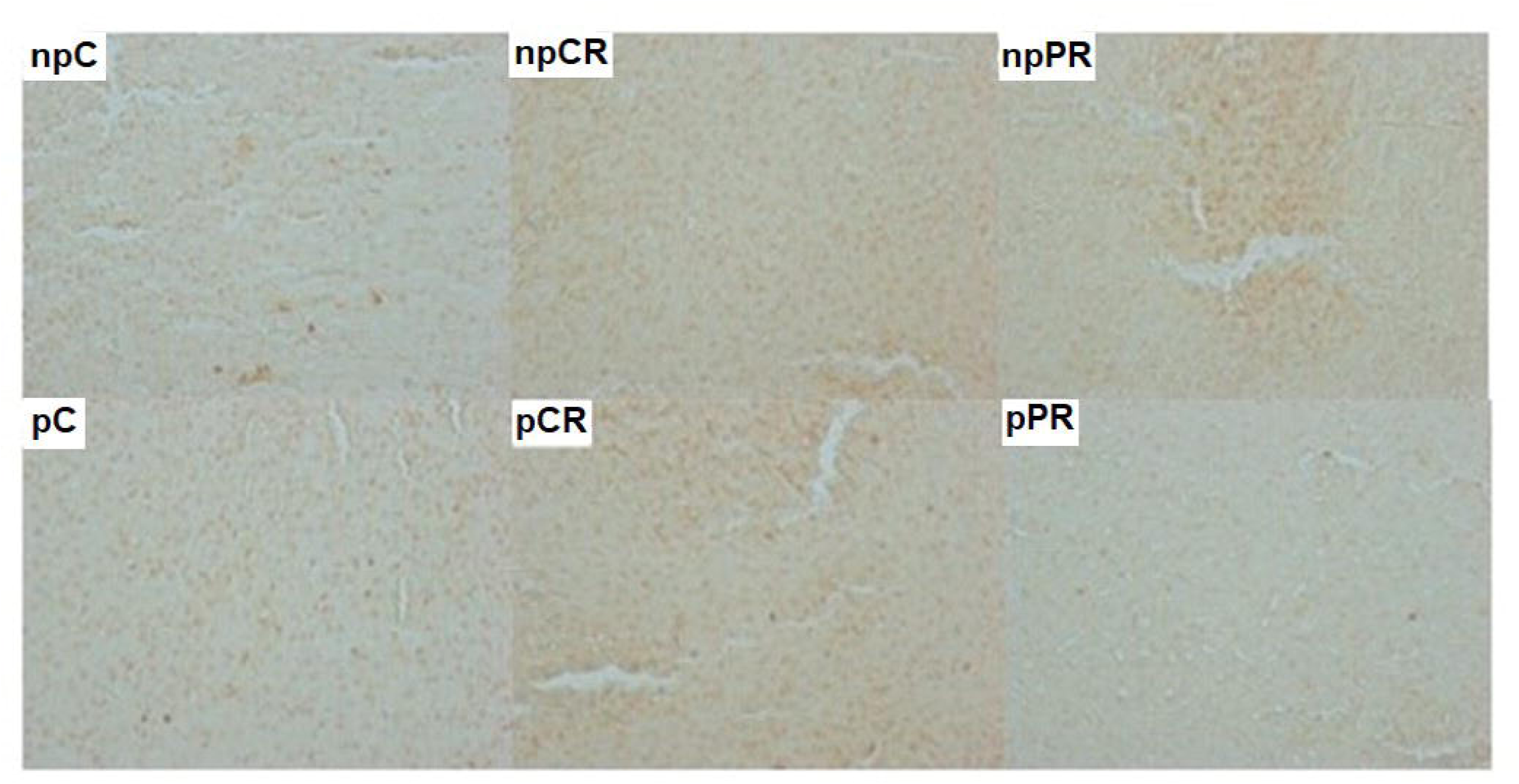
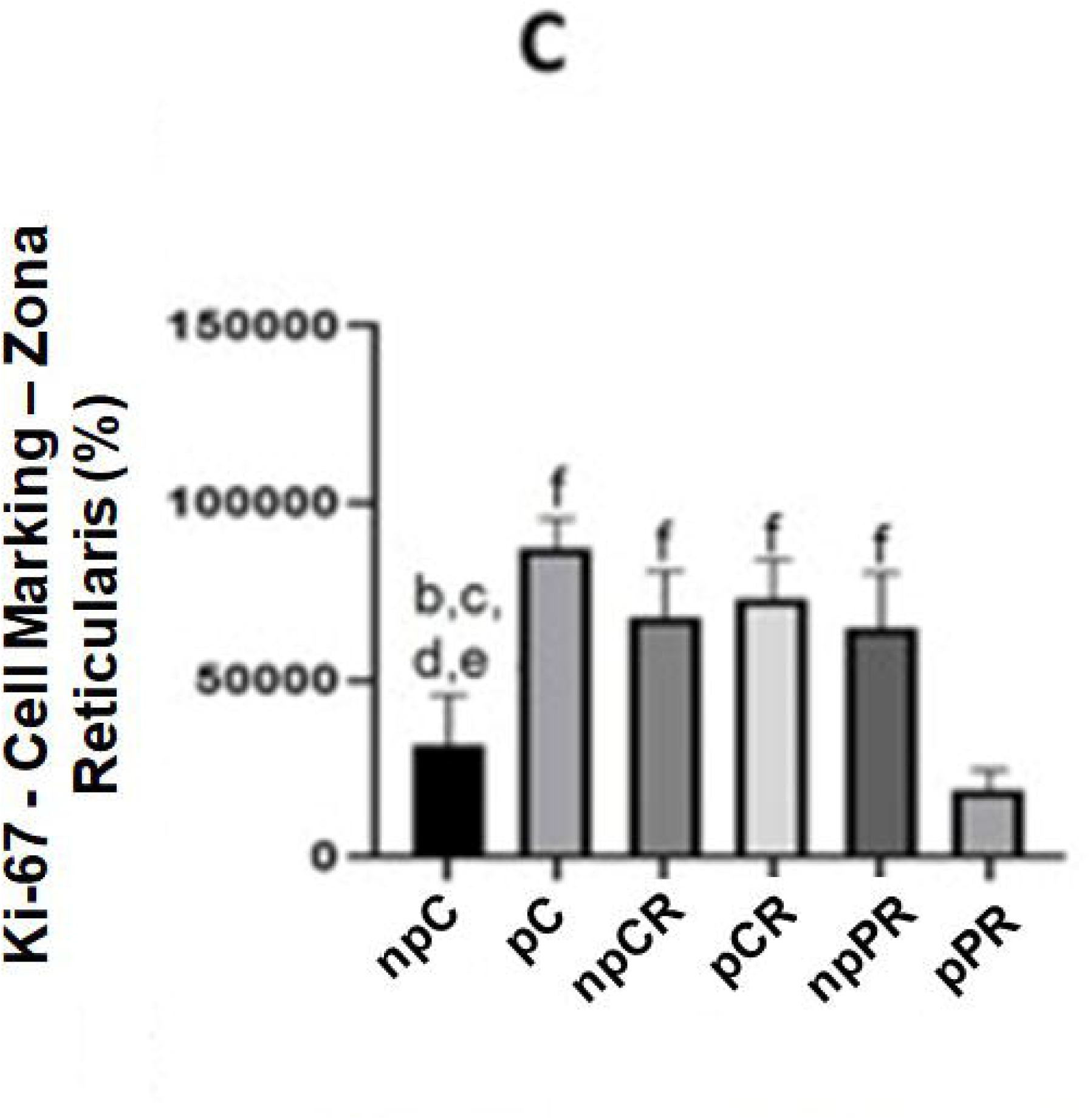
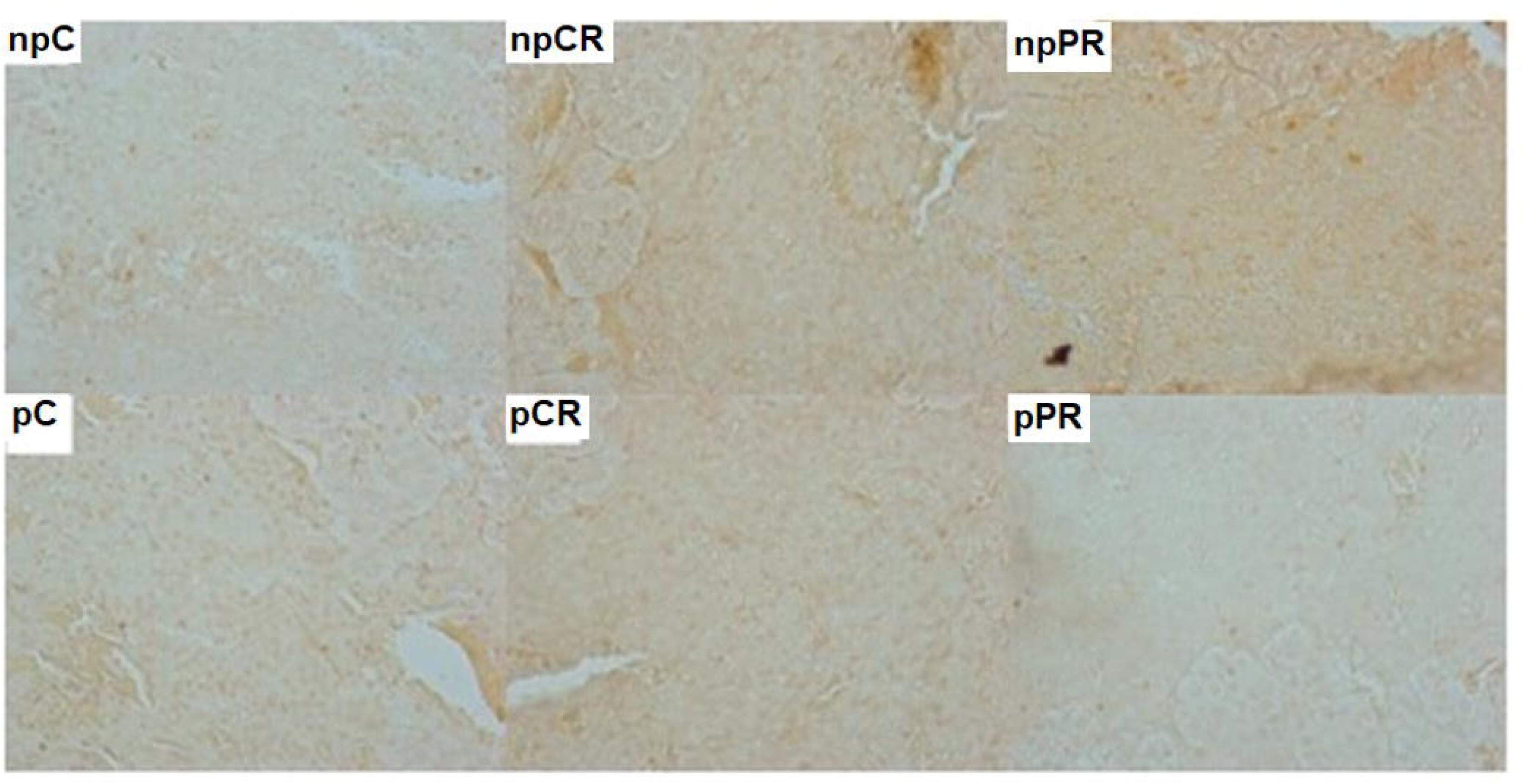
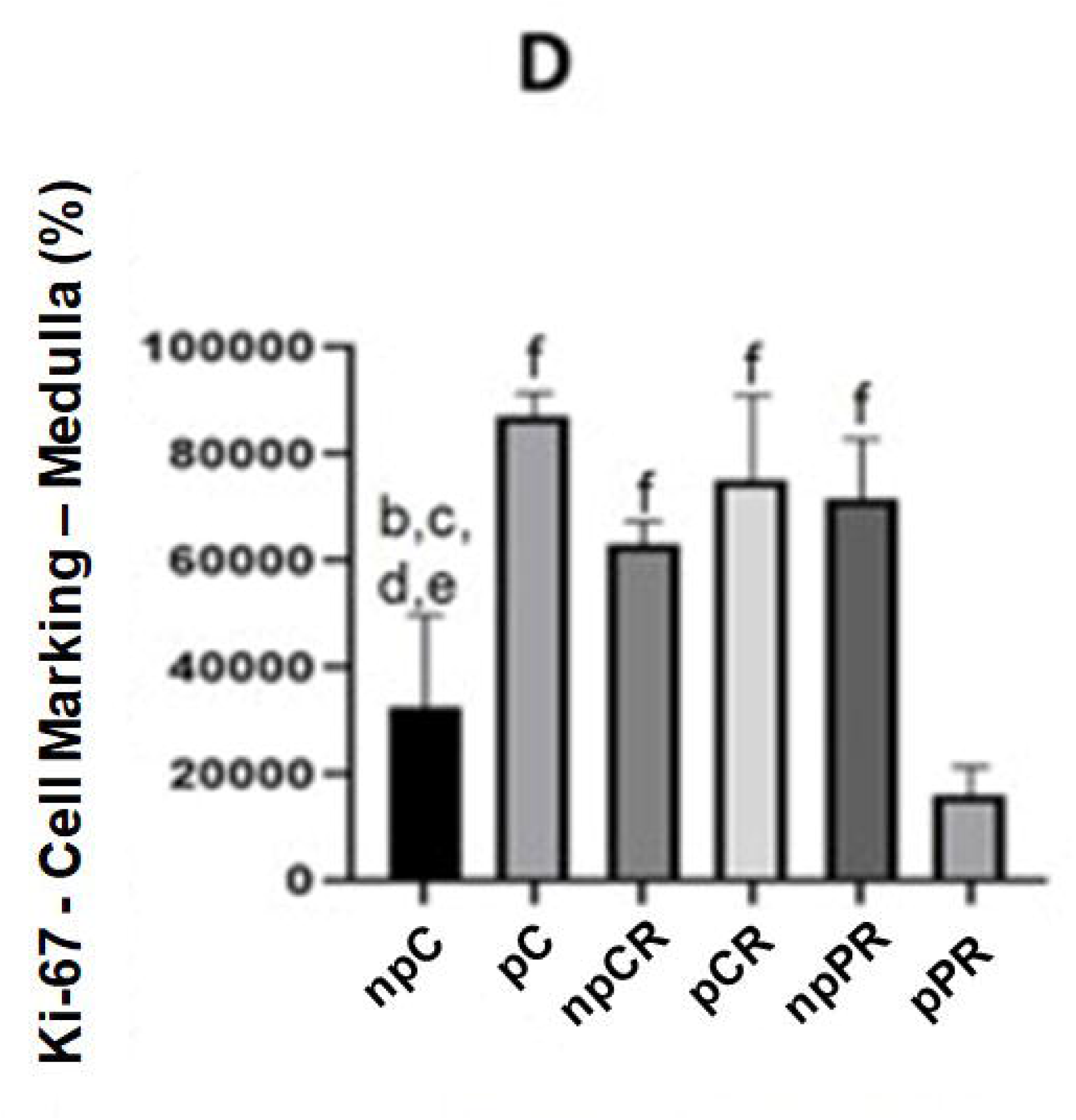
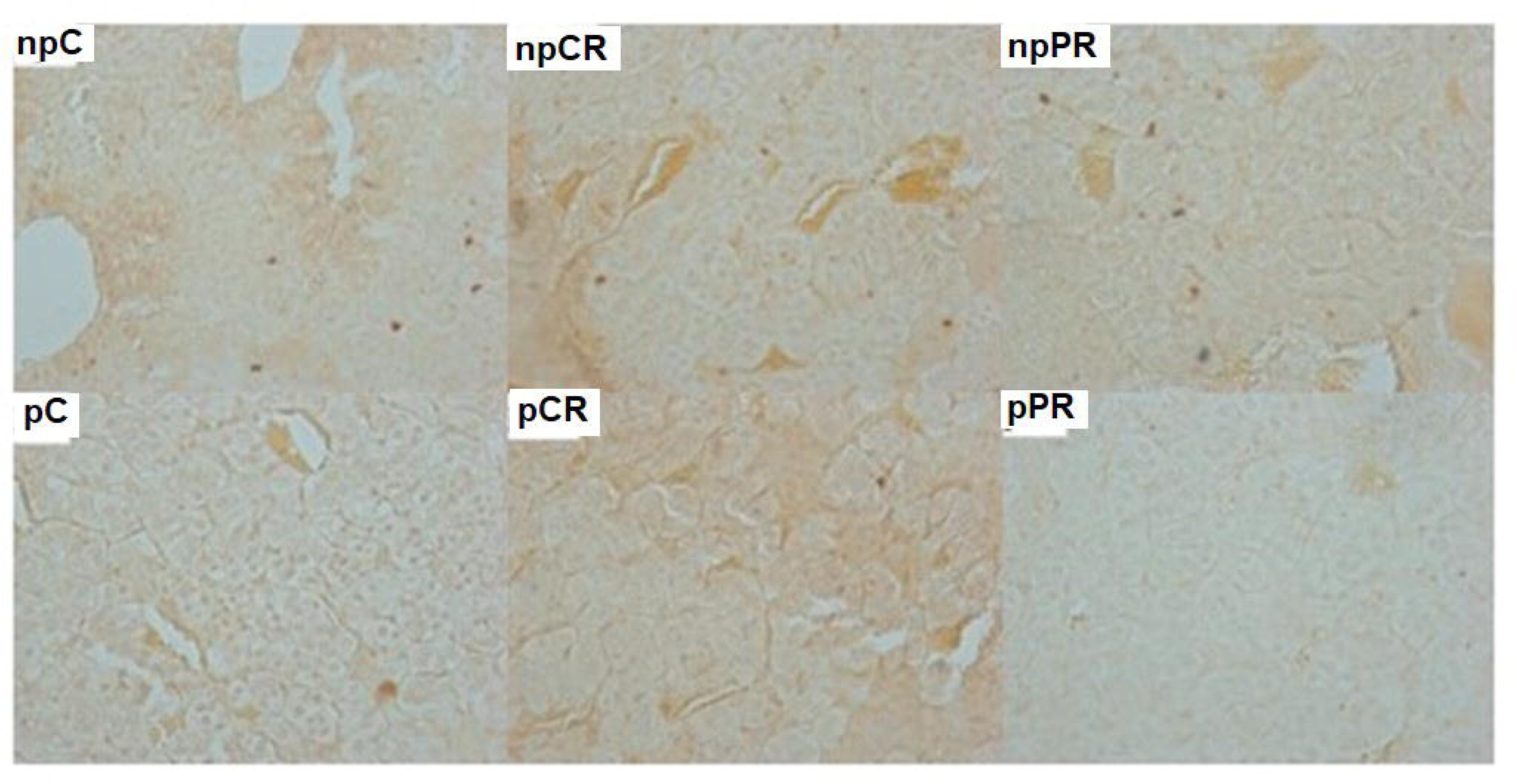
Immunohistochemistry for Ki-67 antigen levels (in percent). Dark brown (stronger markings) and light brown (weaker markings) nucleus markings in their respective areas of the adrenal gland. Final magnification – 200x. A) Glomerular zone. B) Fasciculated zone. C) Reticular zone, D) Medullary zone. Mean ± SD (n = 5; p < 0.05). (ANOVA, post Tukey test).

### 3.5 Effect of diets on GRs in adrenal glands

The zona glomerulosa (Figure 5A) of the npRC group showed the lowest expression of GR compared to those of the npC, pC, and pRC groups. The npRP and pRP groups showed a lower amount of labeling for this receptor than the npC, pC, and pRC groups. GR receptor expression was less marked in the npRP group than in the npC group. The pRP group showed more GR in the zona glomerulosa and fasciculata than the pC and pRC groups (Figure 5).

**FIGURE 5.**
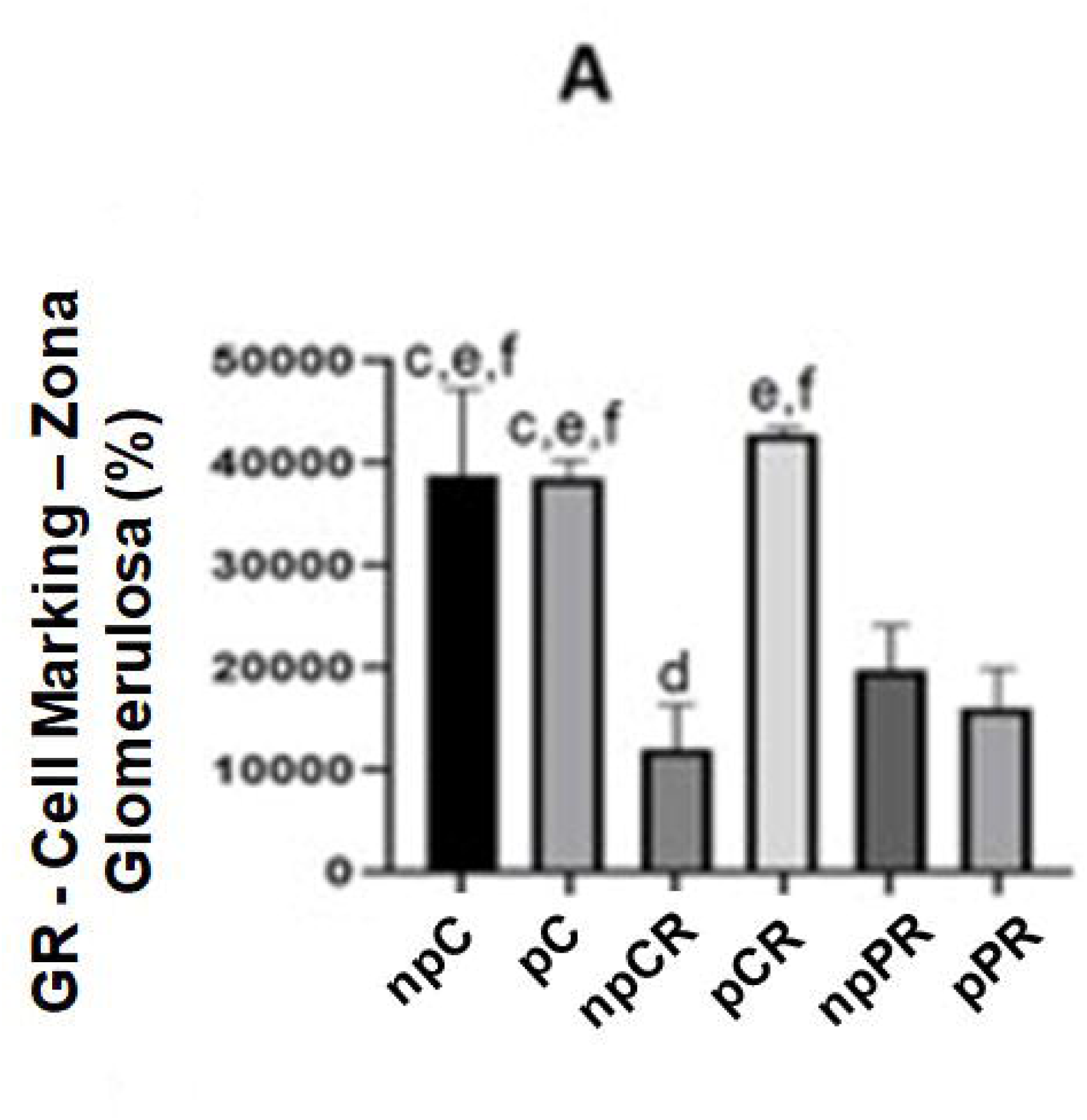
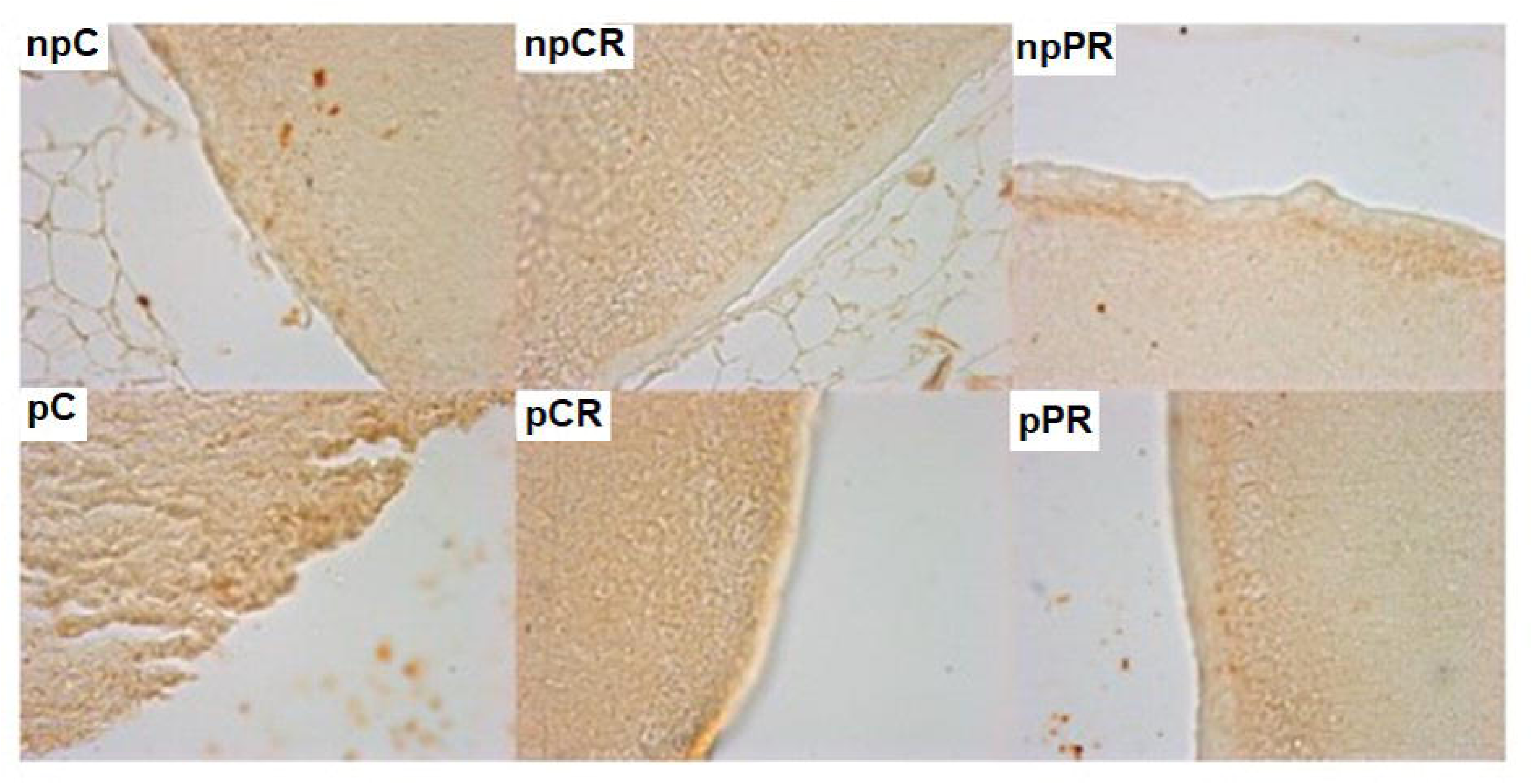
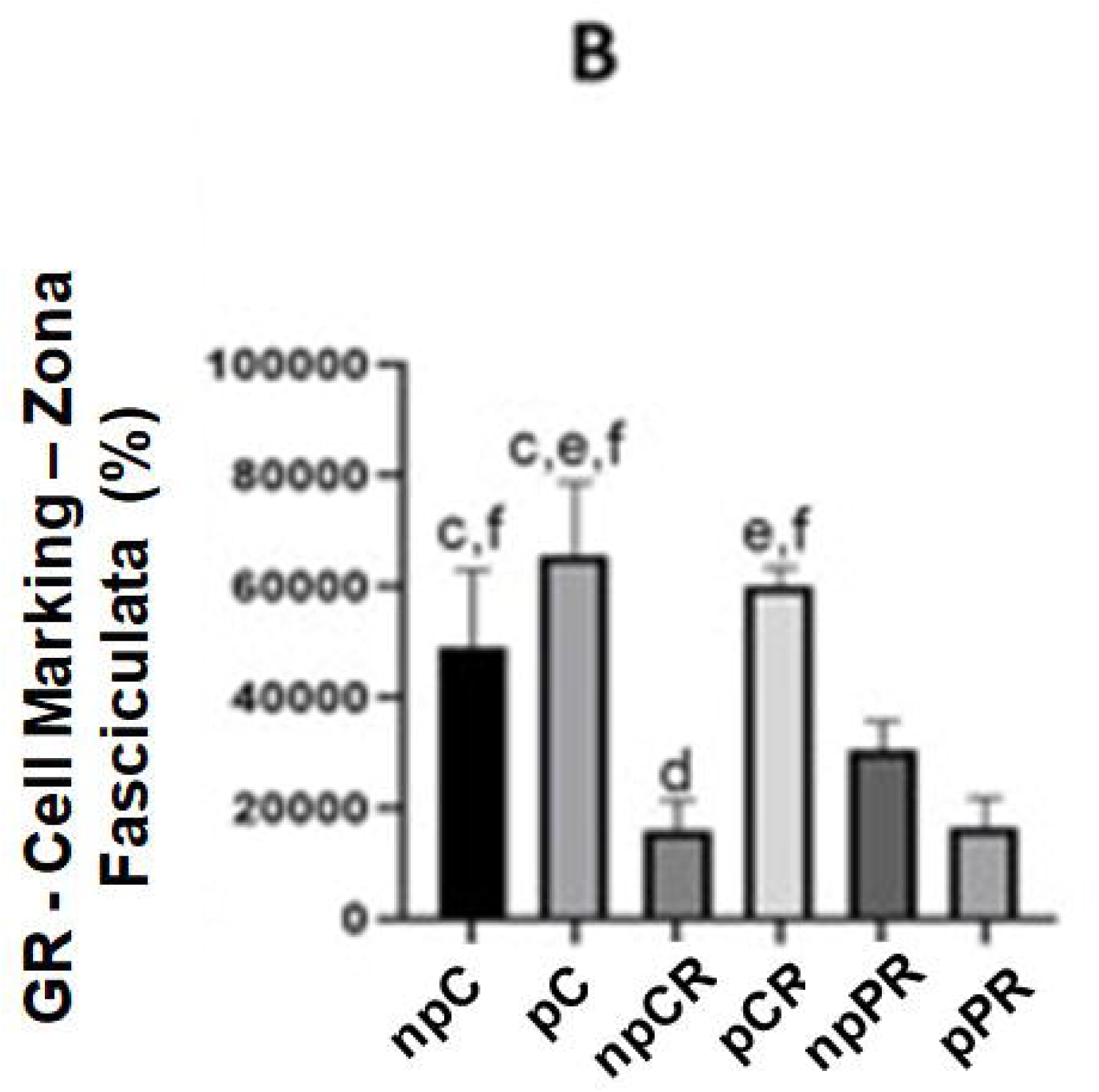
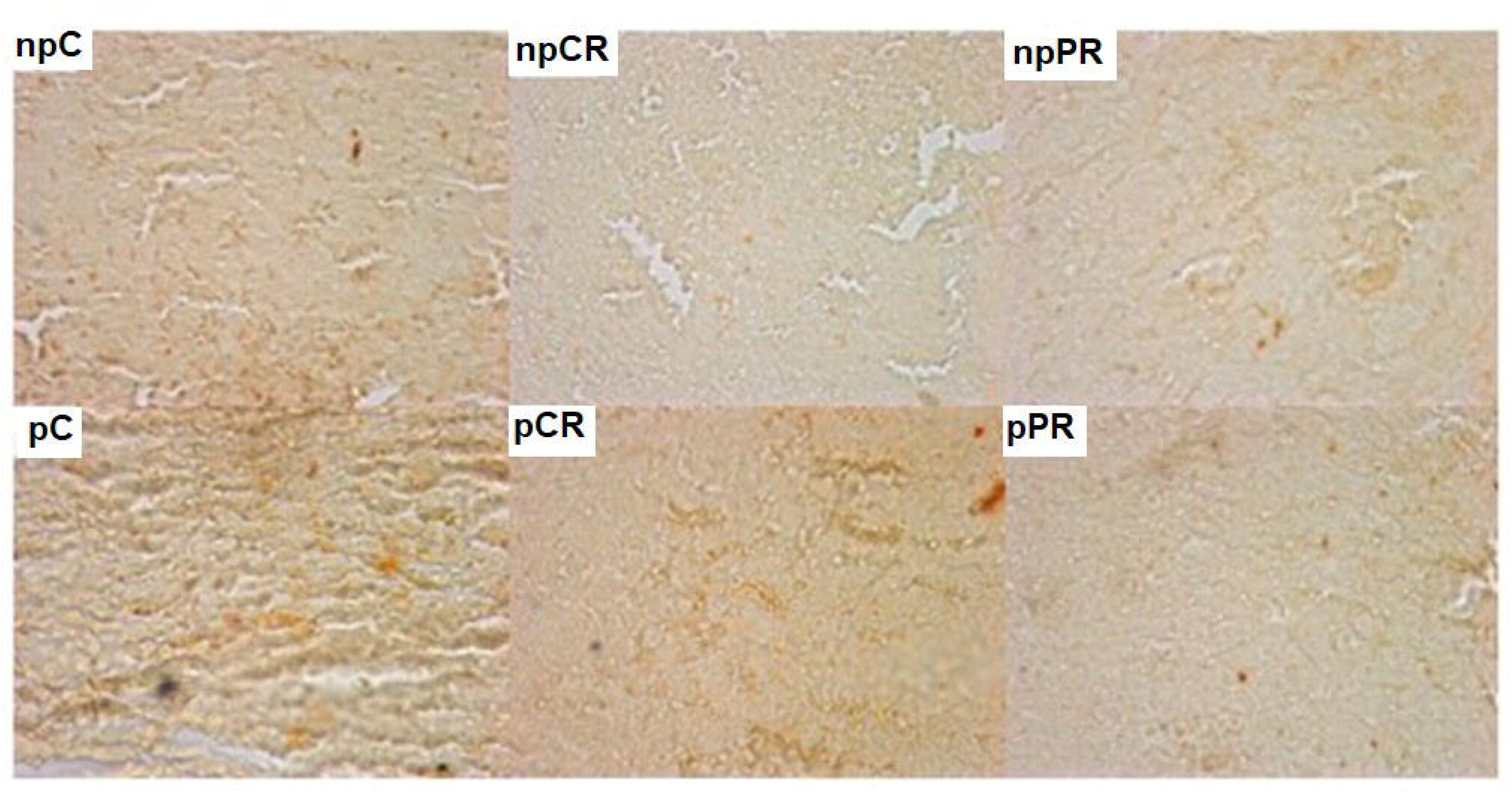
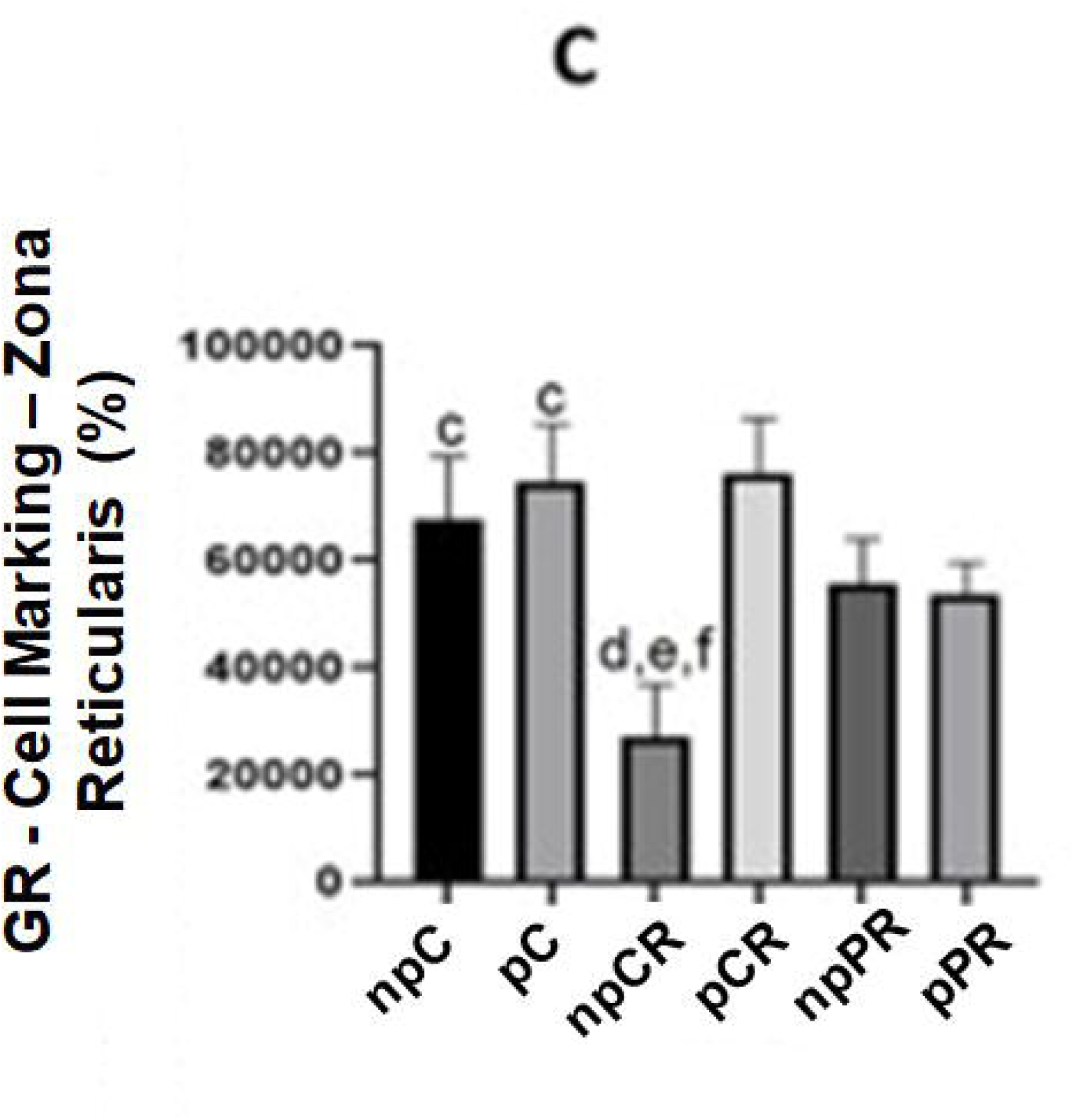
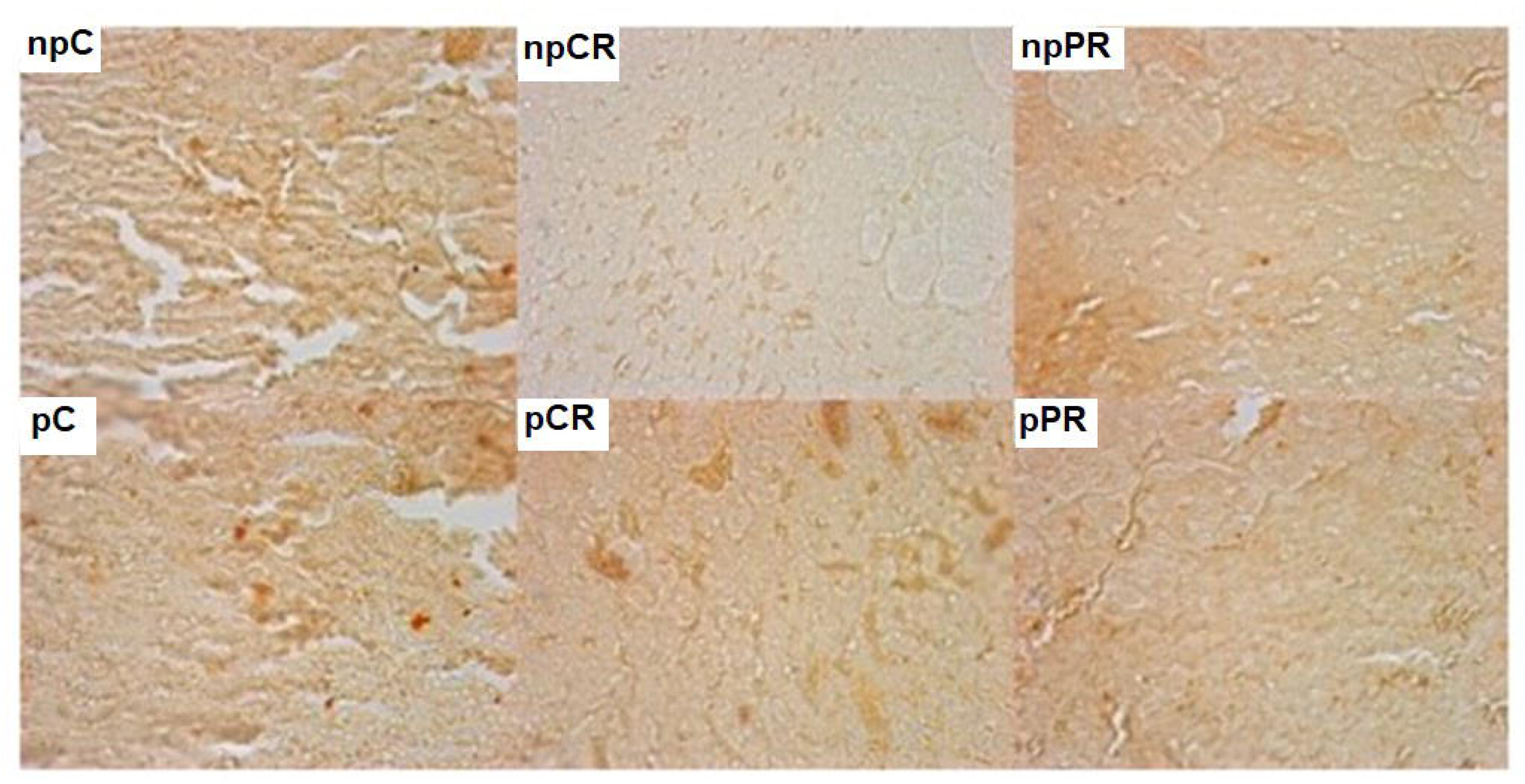
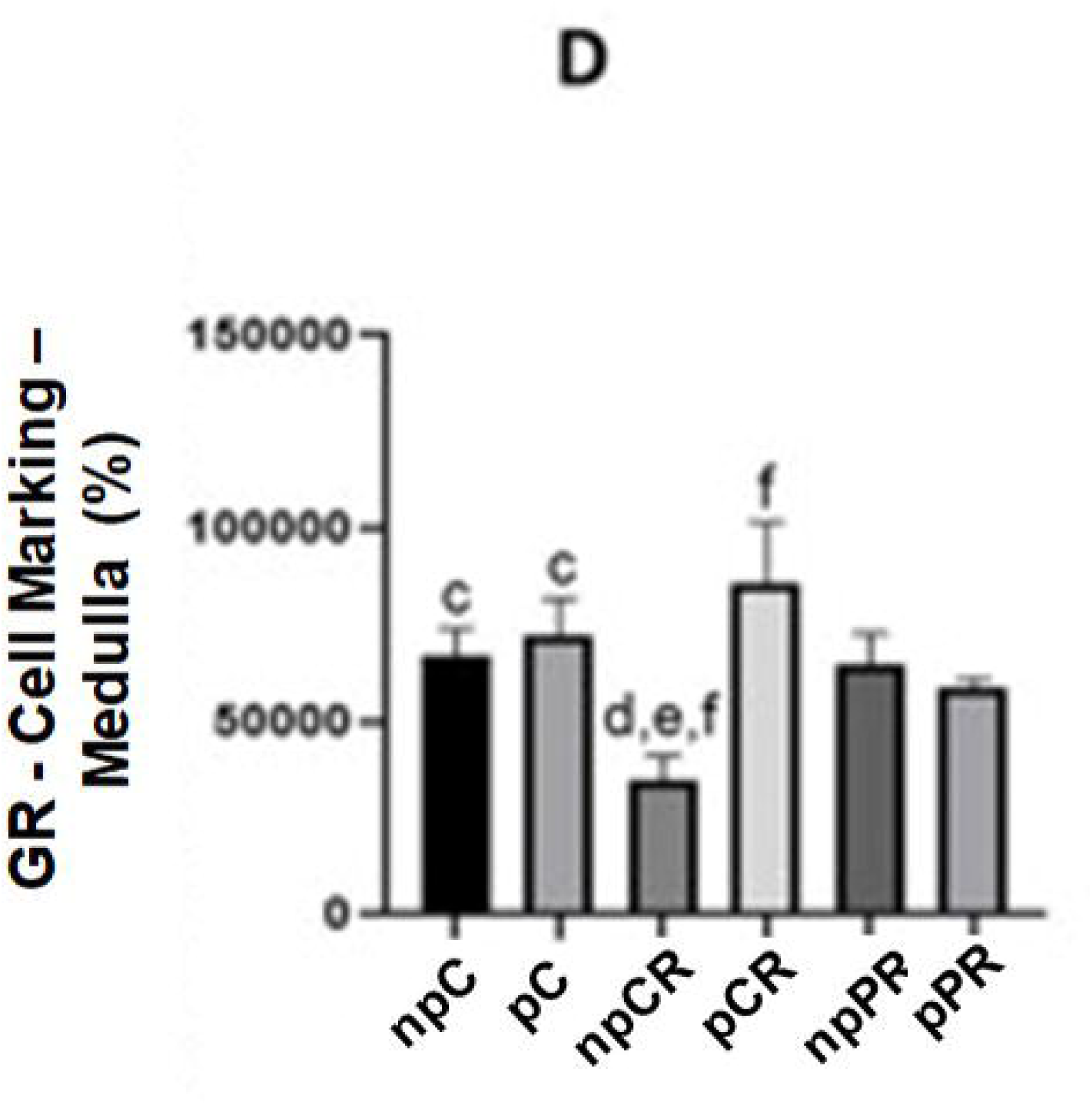
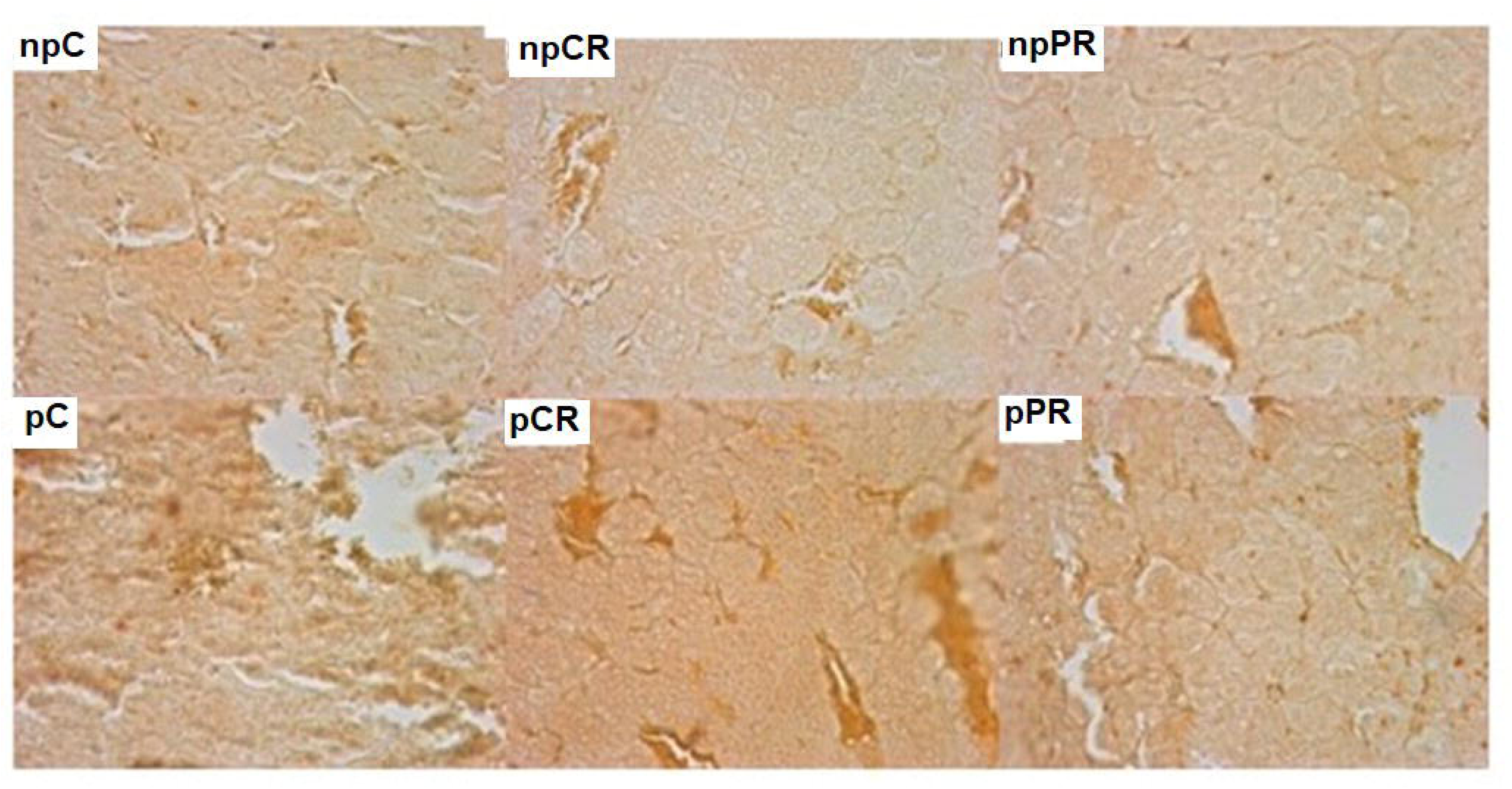
Immunohistochemistry for glucocorticoid receptor (GR) (in percent). Markings in both cytoplasm and nucleus: dark brown (stronger markings) and light brown markings (weaker markings) in their respective areas of the adrenal gland. Final magnification – 200x. A) Glomerular Zone. B) Fasciculate zone. C) Reticular Zone. D) Medullary zone. Mean ± SD (n = 5; p < 0.05). (ANOVA, post Tukey test).

### 3.6 Effect of diet on mineralocorticoid receptors (MR) in the adrenal glands

There were no differences between the control groups in the four adrenal gland zones. However, the pRC group had a greater presence of MR markings than the other five groups. The fasciculate and reticulate zones in the pRP group showed fewer receptors than those in the npRP group (Figure 6).

**FIGURE 6.**
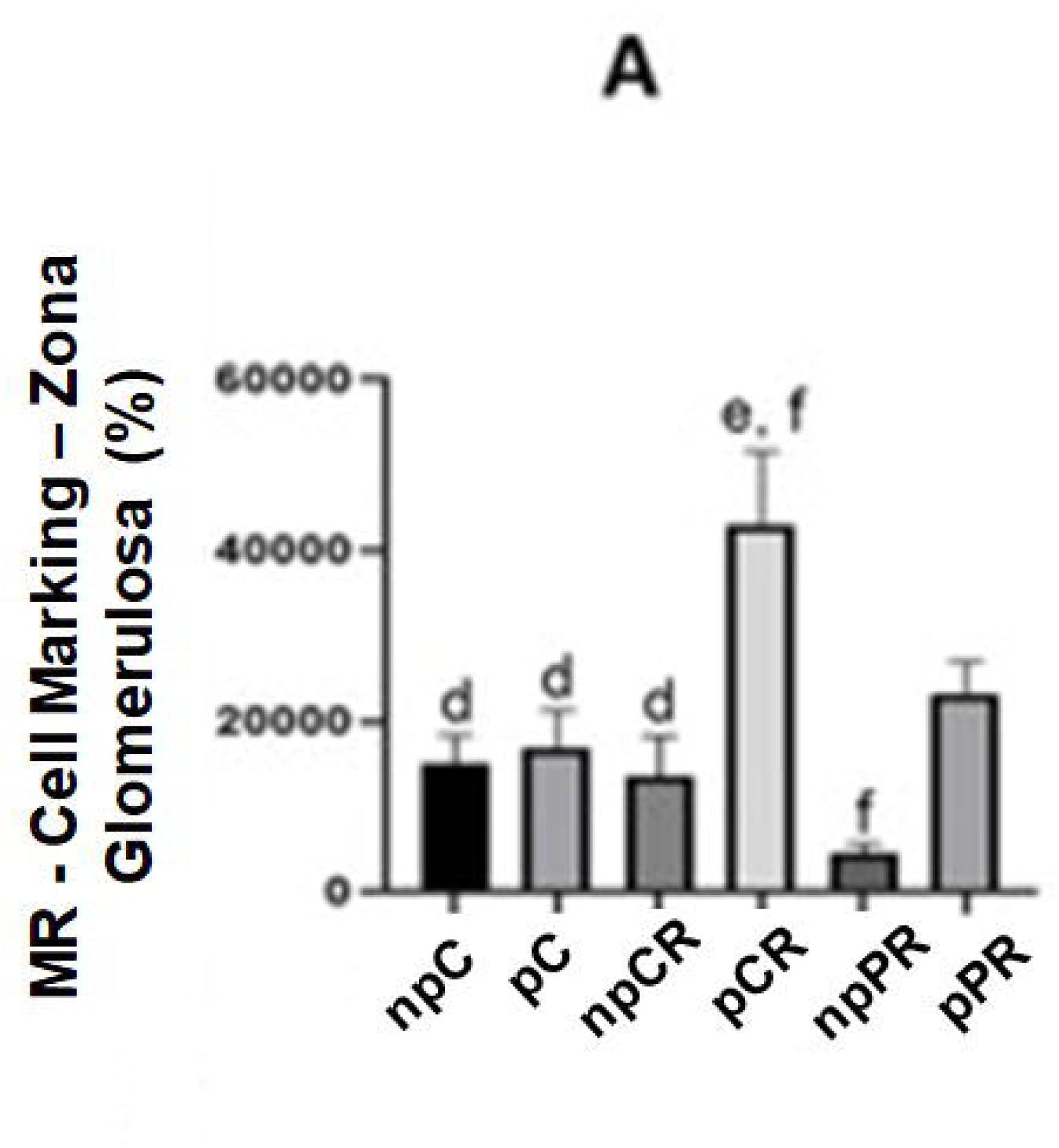
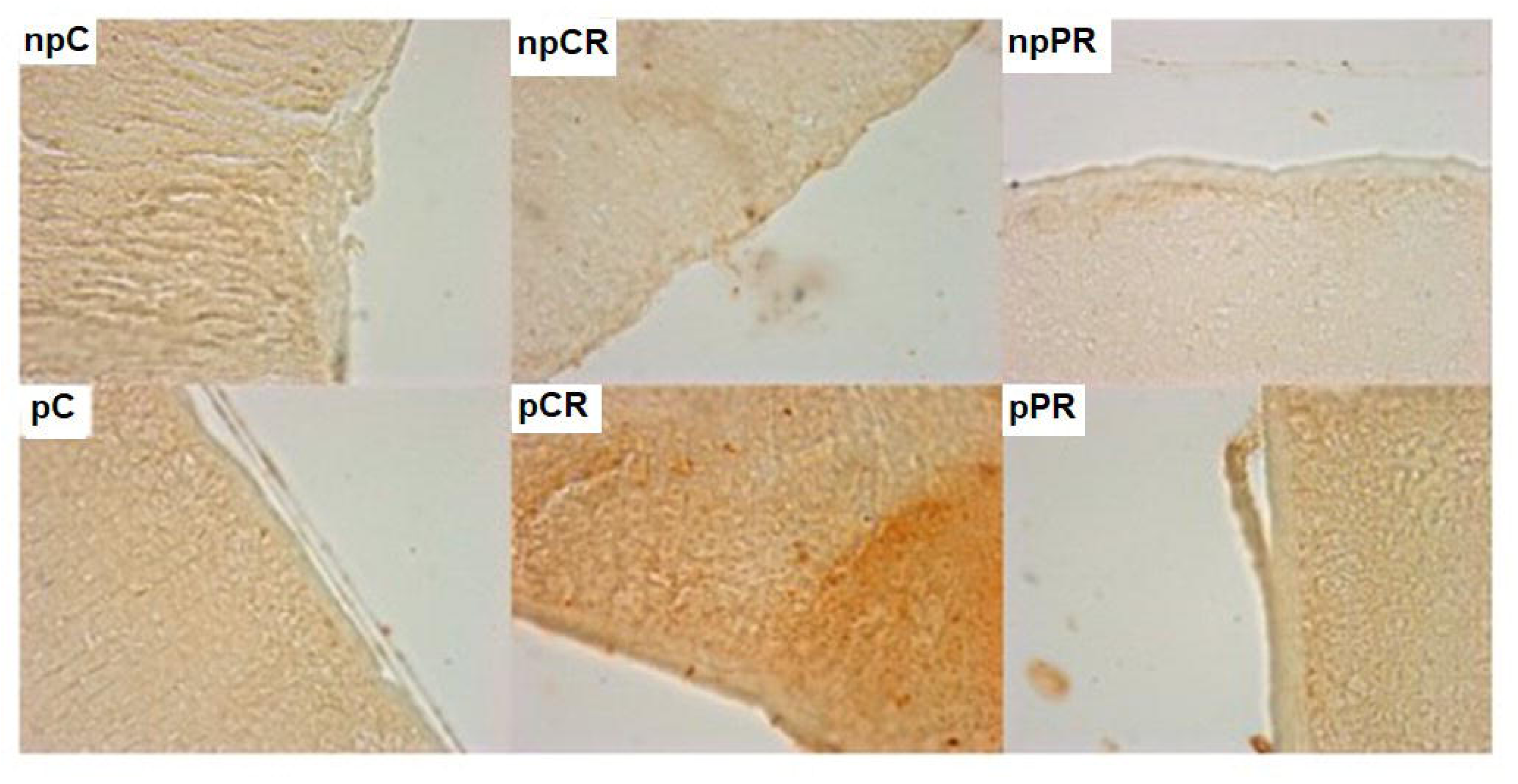
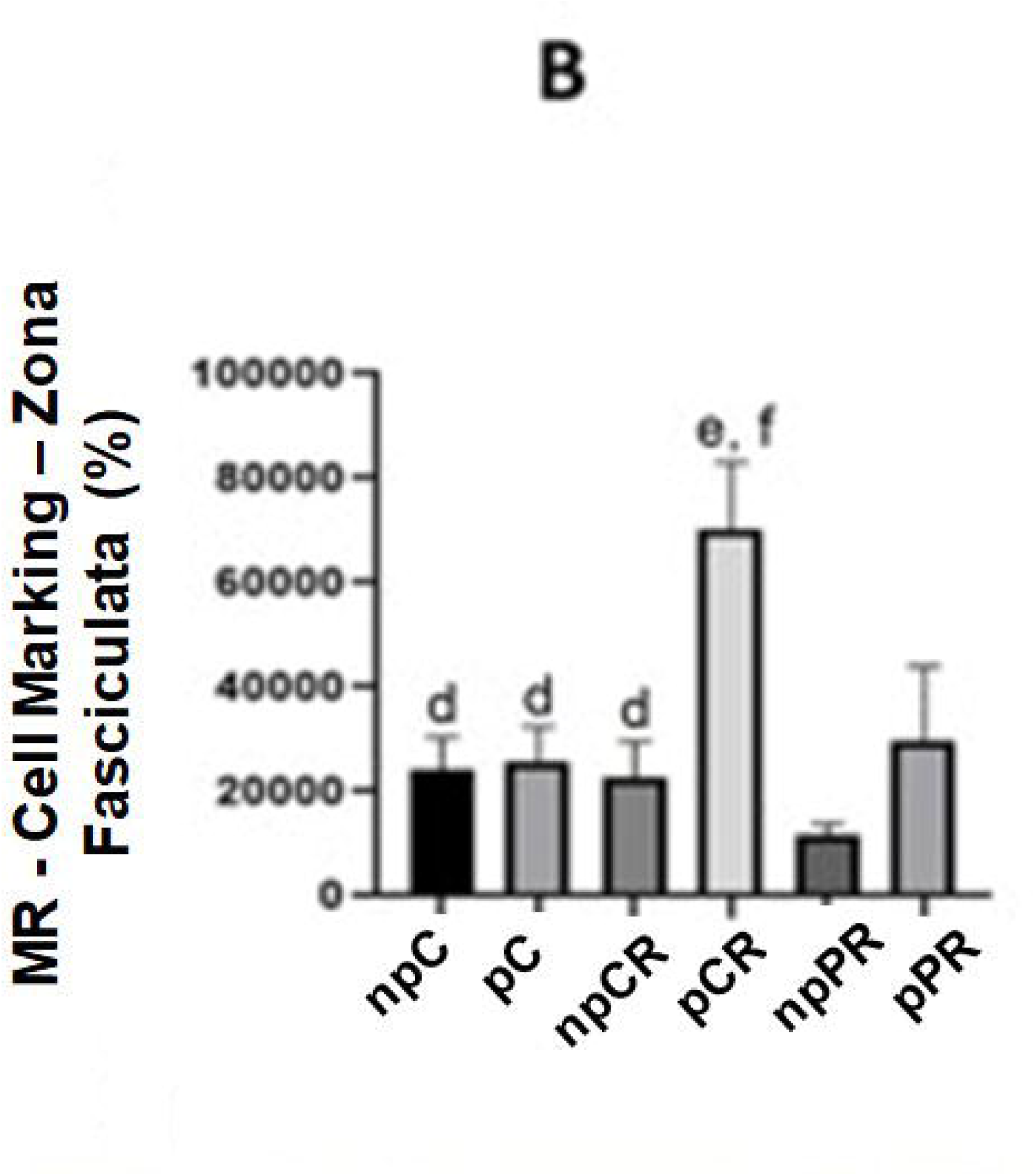
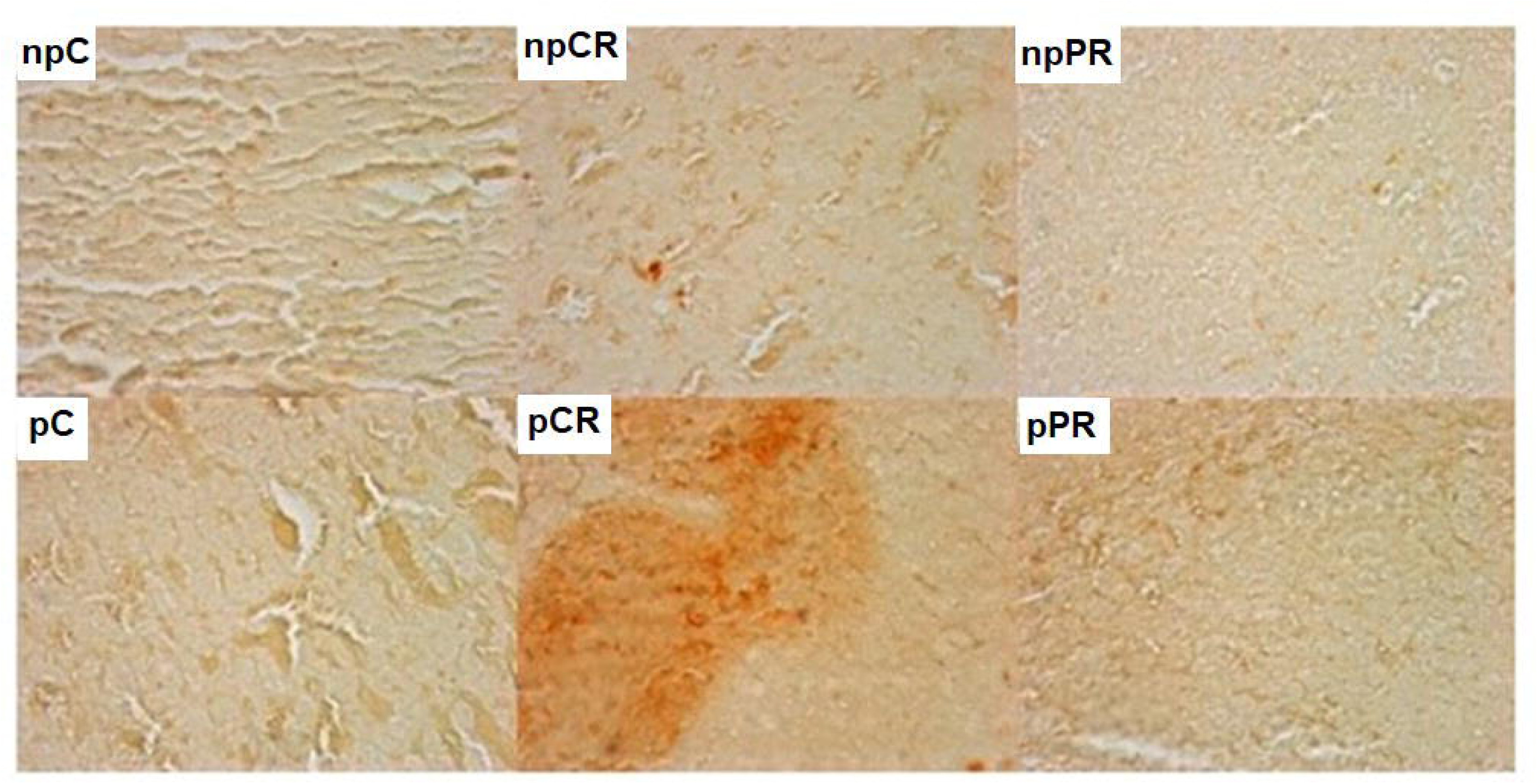
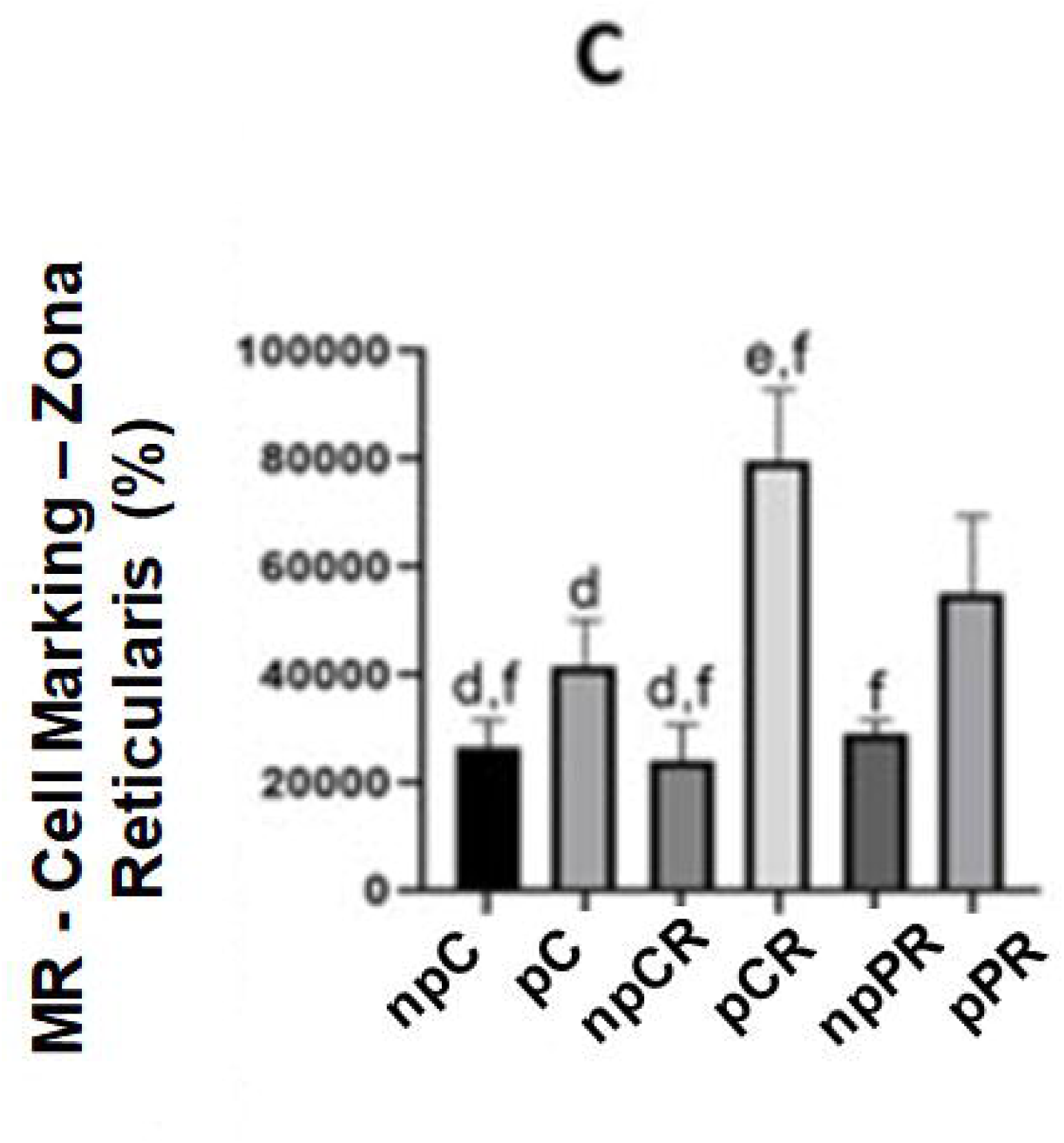
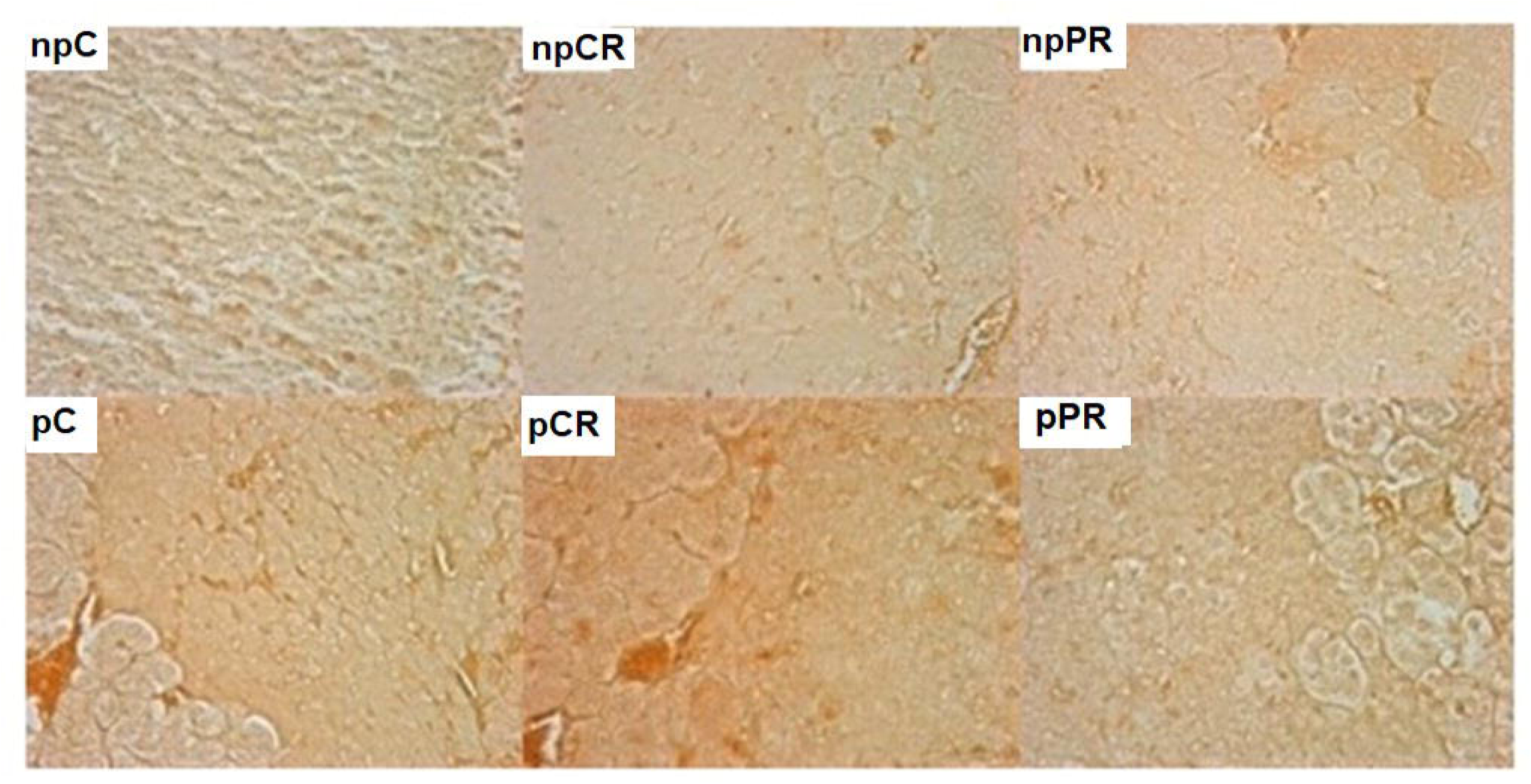
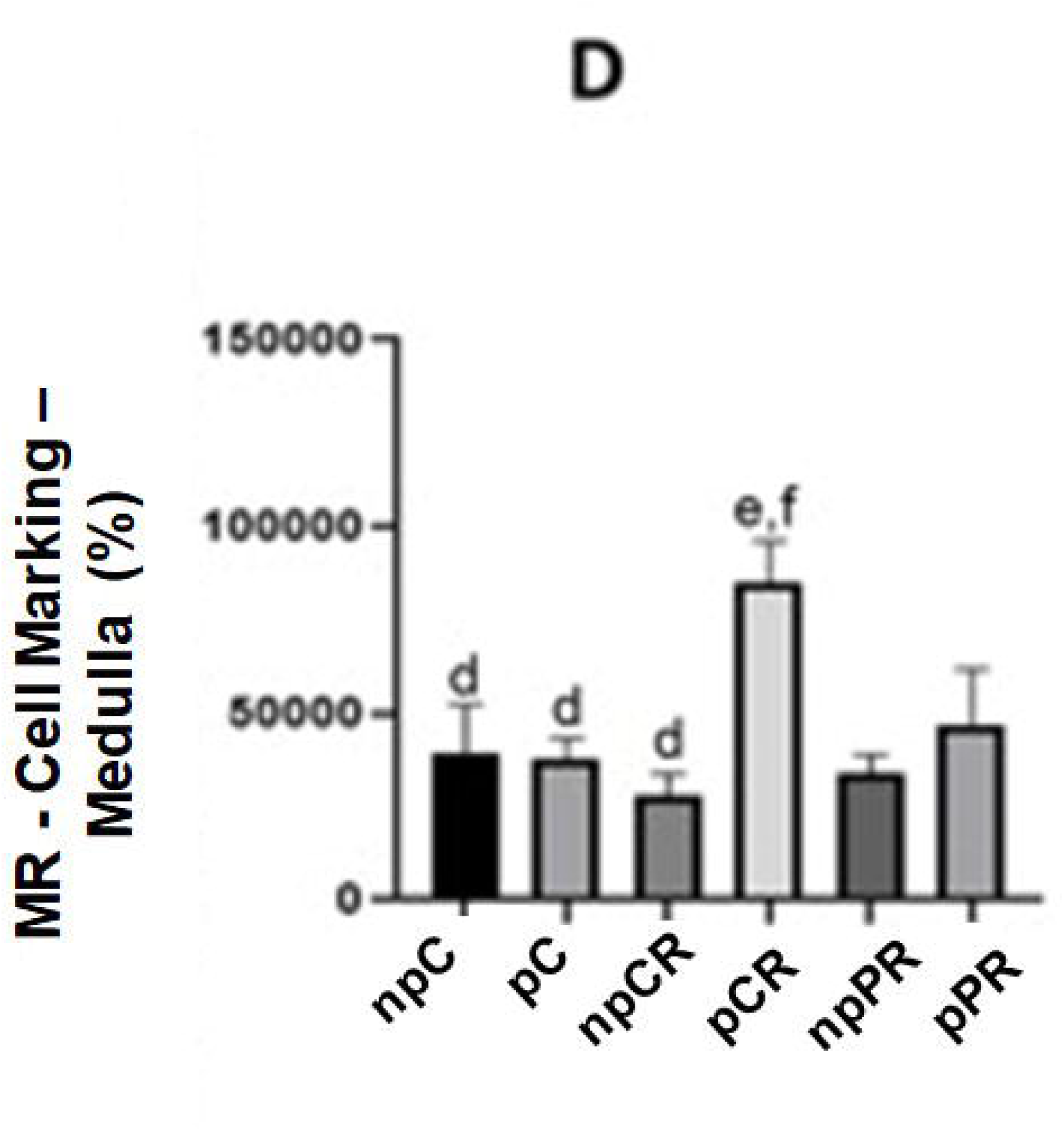
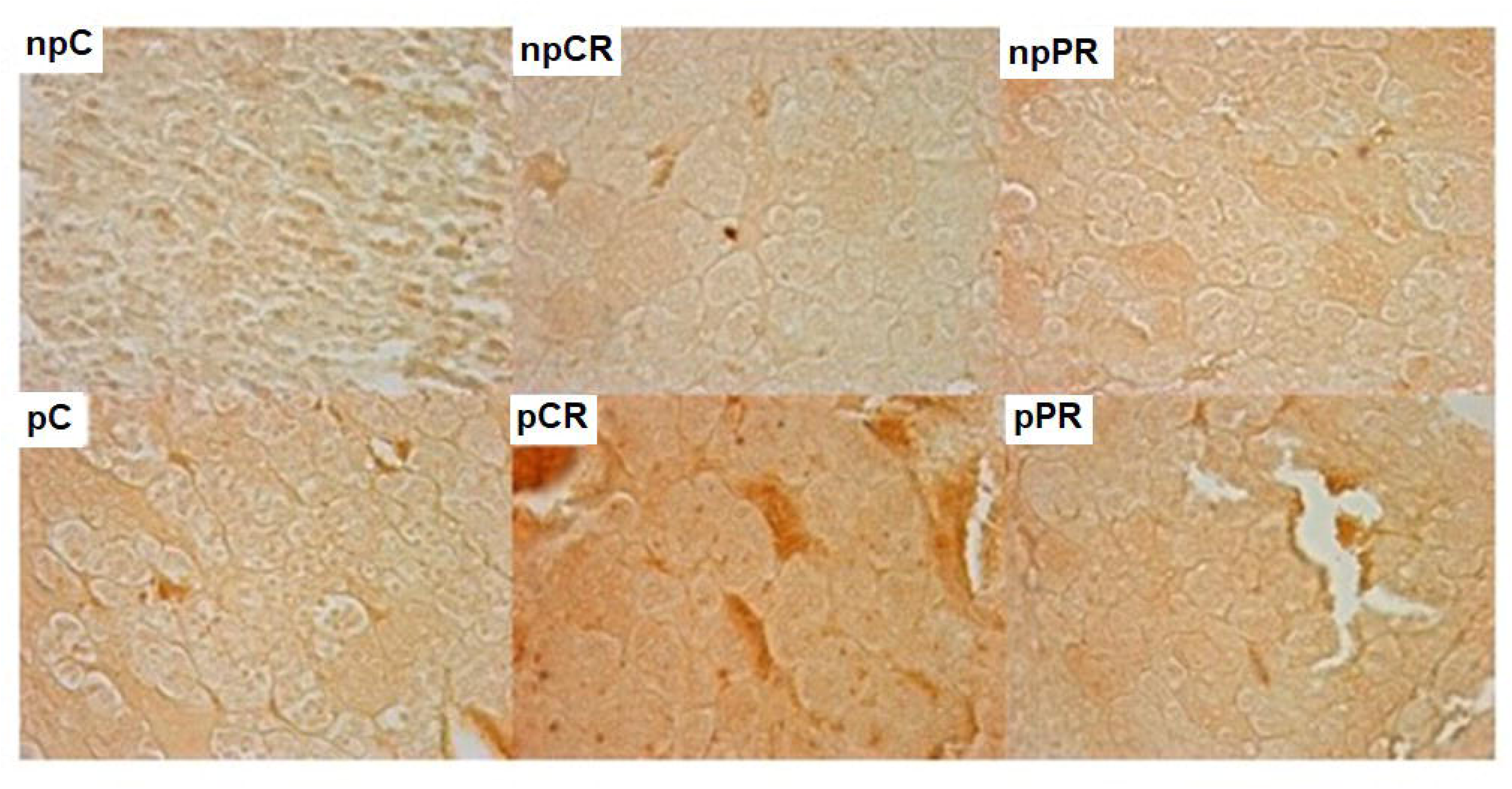
Immunohistochemistry for mineralocorticoid receptor (MR) (in percent). Markings in both cytoplasm and nucleus; dark brown (stronger markings) and light brown (weaker markings) in their respective areas of the adrenal gland. Final magnification – 200x. A) Zona glomerulosa. B) Fasciculated zone. C) Reticular zone. D) Medullary zone. Mean ± SD (n = 5; p < 0.05). (ANOVA, post Tukey test).

## 4 DISCUSSION

In this study, we investigated whether PR and CR during pregnancy could modify the morphology of the adrenal gland as well as the expression of glucocorticoid and mineralocorticoid receptors. Throughout the experimental period, animals presented ascending body mass curves, which indicated normal gestational growth.

CR was validated by the lower food intake of the pRC group during the experimental period. According to the literature, mothers undergoing CR, in addition to having offspring with lower birth weight, also have lower body mass during pregnancy (Barker, 2002). Another finding that validates our study is the upward curve of the pC group, the pregnant group that gained the most mass during pregnancy. A low-protein maternal diet significantly reduced weight gain throughout pregnancy (Cottrell et al., 2012).

Several studies in the literature have demonstrated that the mass of the adrenal gland is restricted, whether in offspring or in mothers, corroborating the data found in this study (Rosenbrock et al., 2005). Huseby et al. (1945) and Boutwell et al. (1948) showed that CR in rats resulted in adrenal hypertrophy without necessarily increasing rat weight (Kritchevsky, 2001). Contrary results have been observed in other studies, in which an increase in adrenal mass in maternal CR of 50% was observed during the last week of pregnancy (Eleftheriades; Creatsas & Nicolaides, 2006) and a smaller adrenal mass was observed (Liang; Zhang & Zhang, 2004).

There have been no specific studies on calorie-restricted or protein-restricted adrenal mass/animal mass ratios. Interestingly, among the nutritionally restricted pregnancy groups, the pRP group had a lower adrenal mass/animal mass ratio than the pRC group. In a study of sodium restriction during pregnancy, there was a 33% increase in the width of the zona glomerulosa in rats. The combination of dietary sodium restriction and pregnancy caused a 167% increase in the width of the zona glomerulosa. There was a direct relationship between the number of cells and the width of the zone, indicating that hyperplasia accompanies hypertrophy of the zona glomerulosa (Pohanka & Pike, 1970), which was not observed in this study, and the cortical and medullary areas did not present significant differences between groups. Future studies are necessary to verify the presence of hyperplasia in a specific zone.

In the presence of inflammation or tissue damage, type I collagen is present in the remodeling of the damaged site (González et al., 2016) and to verify whether restriction in conjunction with pregnancy resulted in increased connective tissue (collagen) deposition in GA, Mallory’s trichrome staining was performed, but no differences were observed between the pregnant and nutritionally restricted groups, which confirms the previous data on the absence of an increase in the thickness of the cortical and medullary areas. Only in the zona glomerulosa, the npC group showed a greater presence of connective tissue (collagen) compared to the pRC groups. Cortisol is known to be responsible for protein degradation (Silverthor, 2010). We noticed, even if discreetly, that the nRC group presented a smaller amount of connective tissue (collagen) in the AG areas, which may indicate that the CR increases stress and consequently cortisol. It has been suggested that a larger RC or RP may present more significant results.

Ki-67 is expressed in cell nuclei during proliferation (Sun & Kaufman, 2018), as a way for tissue to repair damaged cells or meet the organ’s demands. Its expression was higher in the groups with unrestricted pregnancy, with greater cell proliferation in all four areas of the adrenal gland when compared to non-pregnant rats, corroborating the data in the literature (Pohanka & Pike, 1970).

Regarding the zona glomerularis, it was observed that pregnancy with control feeding and with CR had an increase in cell proliferation in the AG in relation to their non-pregnant peers, suggesting possible hyperplasia and hypertrophy in these groups. The opposite occurred in pregnancy with RP, which showed lower cell proliferation than the npRP groups. Although the literature does not report specific studies of KI-67 in RP, some studies on cell proliferation in CR can explain this phenomenon. Studies of this type have indicated a reduction in cell proliferation in keratinocytes, liver cells, mammary epithelial cells, splenic T cells, and prostate cells in 30–50% CR (Bruss et al., 2011; Hsieh et al., 2004; Lok et al., 1990), decreased proliferation of basal cells in the olfactory mucosa of mice (Iwamura et al., 2019), and decreased cell proliferation and an increase in apoptotic cell death (Dunn et al., 1997). The opposite result was observed only in some areas of the brain, such as increased neurogenesis in the dentate gyrus of the hippocampus and the subventricular zone (Kumar et al., 2009; Park et al., 2013; Iwamura et al., 2019). In studies on sheep in the last trimester of pregnancy, the adrenal cortex showed occasional scattered mitotic figures in the zona glomerulosa (Hill et al., 1984).

The only study that addressed the results of cell proliferation in AG was in conducted with a low-sodium diet, where an increase in zona glomerulosa cells was observed together with an increase in aldosterone secretion (Ennen; Levay-Young & Engeland, 2005). Inomata and Sasano (2015) identified greater mitotic activity in the human AG in the region between the zona glomerulosa and zona fasciculata.

Hill et al. (1984) observed that animals subjected to nutritional restrictions, without pregnancy, presented greater proliferative activity in the reticular and medullary zones in relation to the npC group, although in animals fed a normal non-pregnant diet, mitotic activity was absent throughout the adrenal cortex. This finding indicates that RC and RP diets can increase the number of cells in these GA zones.

Elevated levels of Ki-67 and circulating aldosterone expression were associated with treatment with MR antagonists in hypertensive rats, which was observed by immunohistochemistry of ZG cells. This suggests that the greater the hormonal demand by the organism, the more the cells multiplied to supply the hormones (Pereira et al., 2021). This may have occurred in the groups that showed greater cell proliferation, especially the pRC group.

Synthesized or secreted glucocorticoids may play an important role in the direct regulation of adrenocortical cell proliferation and function under physiological conditions (Saito et al., 1979). GR expression in the human adrenal cortex was originally demonstrated by Loose et al. (1980) and was later confirmed in more recent investigations (Paust et al., 2006; Asser et al., 2014). In humans, GR is expressed in the adrenal cortex with functions parallel to those found in other tissues (Briassoulis et al., 2011; Spiga et al., 2017). In addition to its effect on the pituitary and hypothalamus, the cortisol can affect its own synthesis via a local feedback mechanism within the adrenal gland (Gjerstad, Lightman & Spiga, 2018).

MR and GR are present in the adrenal gland, zona glomerulosa, fasciculata and reticulate in humans (Boulkroun et al., 2010), and in sodium-restricted/no MR and GR mice were highly expressed in ZG and ZF/ZR cells (Chong et al., 2017).

GR and MR receptors were observed in newborn rats whose mothers had undergone 50% food restriction during the last week of gestation. Food restriction induces a delay in intrauterine growth, disrupts the HPA axis, and decreases adrenal weight, which was not observed in this study. In addition, newborn mice showed a reduction in MR and GR mRNA in the hippocampus, reduction of CRH mRNA in the paraventricular nuclei of the hypothalamus, and reduction in the plasma levels of adrenocorticotropic hormone (Léonhardt et al., 2002). The same was observed in the hypothalamus of offspring with maternal RP (Bertram et al., 2001), which confirmed our results for GR in the pRP group, which was lower than that in the pC group. The inverse occurred in the expression levels of GR mRNA, which were significantly higher in the kidney, lung, liver, and hippocampus of fetal and neonatal pups at the end of gestation (20 days) and in 12-week offspring exposed to maternal RP, suggesting that this increase is persistent throughout life (Bertram et al., 2001). MR labeling showed that in the zona glomerulus and reticularis, there was greater labeling in the pRP group than in the npRP group, which suggests that pregnancy influences the increase in MR, with glucocorticoids activating MR in most tissues at baseline levels and the GR at stress levels (Gomez-Sanchez & Gomez-Sanchez, 2014).

GR mRNA expression levels in nutrient-restricted neonatal ewe pups were the highest in adrenal, kidney, liver, lung, and perirenal adipose tissues, where the persistence of tissue-specific increases in GR, 11β-hydroxysteroid dehydrogenase type, was demonstrated. 1-11bHSD1 (which transforms cortisone to cortisol) and angiotensin II receptor type 1 (AT1) decrease the expression of 11β-hydroxysteroid dehydrogenase type 2-11bHSD2pa (which oxidizes cortisol) in the adrenals and kidneys of newborn infants in response to a defined period of maternal nutrient restriction during early pregnancy. The authors inferred that gene expression is programmed by the availability of nutrients to the fetus before birth (Bertram et al., 2001). This may explain the large increase in MR and GR receptors in the pRC group; due to food restriction, there was a greater production of cortisol and stress due to the greater expression of MR and GR.

Malnutrition in pregnant rats in GA causes a decrease in GR mRNA expression due to stress, which increases the maternal production of corticosterone. This makes tissues most sensitive to corticosteroid concentration. This in turn stimulates The expression of HSD11B1 in rats (Khorram et al ., 2011) causing excess maternal and fetal plasma corticosterone, downregulating fetal GR and MR, and compromising the HPA feedback axis in childhood and adulthood (Valsamakis, Chrousos & Mastorakos, 2019). Although the groups did not suffer from malnutrition in the glomerular, fasciculate, and medullary zones, the pRC group showed greater staining for GR when compared to the pRP group, and the same was repeated for the MR staining in the four AG zones. The results suggest that the groups may have experienced stress of restriction, with CR having an impact on rats during pregnancy compared to RP. However, the results were different from those found in the literature since there was no negative regulation of the receptors, as the expression levels of MR and GR were higher in the pRC group.

The negative regulation of the receptors was noted by lower GR labeling in the npRC group than in the npC group. Although there are no comparative studies between PR and CR in the literature, we present in this study that in the reticular and medullary zones, the npRC group had lower markings for GR compared to the npRP group, and the opposite occurred in the medullary zone when compared to the CR pregnant groups and the PR groups. MR expression levels were unaffected by the maternal diet in the kidneys of offspring of maternal RP and were undetectable in the lungs (Bertram et al., 2001).

In all GA, both GR and MR were expressed more in pRC than in npRC, suggesting that pregnancy is an additional stressor because in pregnant primates, there was an increase in maternal cortisol (Recabarren, Valenzuela & Seron-Ferrer, 1997). However, in a study in adult male rats with perinatal malnutrition, the levels of GR and MR mRNA expression and the binding capacity or affinity showed no difference between groups in GA (Dutriez-Casteloot et al., 2008).

In the offspring of pregnant hamsters, the rate of steroidogenesis increased in malnourished rats (Liang, Zhang & Zhang, 2004) and although we did not measure cortisol directly, the results of the pregnant group with CR showed an increase in the number of GR and MR receptors, increased cell proliferation, and indirect features for an increase in cortisol production by GA cells.

It has been suggested that aldosterone secretion in the rat ZG can be regulated by MR through ultra-short *feedback* within the adrenal gland, where aldosterone regulates its own production, and because it can be activated by the same hormone, GR can also regulate the production of glucocorticoids in the ZF and ZR (Chong et al., 2017). This phenomenon may explain our results on the increase in the number of receptors that may regulate cortisol production without necessarily increasing the number of cells for this role.

However, there is disagreement as to whether the *feedback* exerted by the MR/GR receptors within the adrenal gland is positive or negative. Some studies have stated that the *feedback* provided is positive. *In vitro* research with H295R cells (a human cortisol-secreting adrenocortical cell line) revealed the presence of an intra-adrenal positive feedback loop that regulates steroid production. These results were confirmed when GR was inactivated by the pharmacological antagonist RU486 or GR *knockdown* by siRNA, which led to the suppression of steroidogenesis, strongly suggesting an autocrine and GR-mediated ultra-short autocrine positive regulatory loop (Asser et al., 2014). This type of feedback corresponds to the results of the pRC groups, in which there were increases in GR and MR receptors in response to past stress during the pregnancy and CR periods.

Chong et al. (2017) pointed out that this regulation occurs through negative feedback because MR and GR negatively regulate glucocorticoid production in ZF/ZR cells (intra-adrenal feedback–short loop). *In vitro* and *in vivo* studies have shown that prior exposure of the adrenal gland to glucocorticoids results in a diminished response to adrenocorticotropic hormone (ACTH), resulting from an intra-adrenal negative *feedback loop* that could constitute an additional GR-regulated control mechanism for steroidogenesis (Peron et al., 1960; Carsia & Malamed, 1979; Chong et al., 2017; Gjerstad, Lightman, & Spiga, 2018). This may justify that even with an increase in cell replication, increased measurements in the adrenal zones in some groups are not sufficient to increase cortisol production as they are regulated by negative intra-adrenal *feedback* by MR and GR.

The results showed that PR and CR during pregnancy did not change the weight of the adrenal glands when compared to non-pregnant women. RC increased the expression of GR and MR receptors in GA during pregnancy, whereas RP decreased the labeling of GR and MR in the zona glomerulosa and fasciculata. We concluded that CR during pregnancy caused the most stress to the rats, altering the presence of MR and GR, which may suggest an alteration in the functionality of the GA and, consequently, in the HPA axis.

## ACKNOWLEDGMENTS

This research was supported by the Hermínio Omettto Foundation.

## AUTHOR CONTRIBUTIONS

**Bruno dos Santos Telles**: Carried out experimental work, data collection and data evaluation. **Hércules Jonas Rebelato**: Carried out experimental work, data collection and data evaluation. **Marcelo Augusto Marretto Esquisatto**: Conceptualization, methodology, validation, formal analysis, investigation and writing. **Rosana Catisti**: Conceptualization, methodology, validation, formal analysis, investigation, writing - review & editing of manuscript and supervision.

